# Spatial patterns of phylogenetic diversity and endemism in the Western Ghats, India: a case study using ancient predatory arthropods

**DOI:** 10.1101/2020.10.19.344796

**Authors:** D. K. Bharti, Gregory D. Edgecombe, K. Praveen Karanth, Jahnavi Joshi

## Abstract

**Aim:** To study patterns of phylogenetic diversity, endemism and turnover in a community of ancient arthropods across a biodiversity hotspot. Our specific aims were to understand diversity gradients, identify hotspots of endemism and conservation importance, and highlight poorly-studied areas with unique biodiversity.

**Location:** The Western Ghats (WG), India

**Methods:** We compiled a location data-set for 19 scolopendrid centipedes species which was used to predict areas of habitat suitability using bioclimatic and geomorphological variables in Maxent. We used predicted distributions and time-calibrated species phylogeny to calculate taxonomic and phylogenetic indices of diversity, endemism and turnover.

**Results:** We observed a decreasing gradient in Taxonomic and Phylogenetic Diversity (TD/PD) from the southern to northern WG and high Phylogenetic Endemism (PE) in the southern and northern WG. Southern WG had the highest diversity and was represented by lineages with long branch lengths and short ranges as observed from Relative Phylogenetic Diversity/Endemism (RPD and RPE). Despite having low PD, the northern WG had high values of PE represented by distinct lineages as inferred from RPE. Sites across the Palghat Gap grouped separately in comparisons of species turnover along the WG.

**Main conclusions:** Our findings support expectations from the latitudinal diversity gradient in the WG and the southern WG refuge hypotheses. The high diversity and endemism along with the presence of ancient lineages in the southern WG is consistent with *in-situ* speciation. Climatic differences or dispersal barriers might have retained this diversity locally. High phylogenetic endemism in lateritic plateaus of the northern WG, albeit with low phylogenetic diversity, indicates the presence of distinct evolutionary lineages that might be adapted to life in these landscapes characterized by poor soil conditions and seasonal ephemeral habitats. Our results from soil arthropods highlight the need to use phylogeny and distribution data while assessing diversity and endemism patterns in the WG.

## Introduction

Understanding patterns of species diversity and its drivers remains a challenge in community ecology. There are many ways to characterise a community, though one of the most commonly used measures is taxonomic diversity (TD). Taxonomic diversity treats all species as evolutionarily independent units, but this may not be true. To address this, phylogenetic diversity (PD) was proposed to explicitly incorporate the evolutionary history for each species, which would reflect the accumulated evolutionary history of a community (Faith, 1992). Further, the geographical distribution of a species along with phylogenetic divergence can be incorporated for all taxa in a given region through phylogenetic endemism (PE), a metric that weights the branch lengths of each lineage by its respective geographic range (Rosauer et al., 2009). The use of PD and PE allows us to assess the roles of ecological, historical and evolutionary processes that structure communities, and their usefulness has been demonstrated in multiple biodiverse and complex landscapes (Azevedo et al., 2020; Fenker et al., 2020; Mishler et al., 2014), but remains limited in the Asian tropical forests where only PD has been assessed (Bose et al., 2019; Divya, Ramesh & Karanth., 2020; Tamma & Ramakrishnan, 2015).

One of the limiting factors for assessment of species diversity and endemism over large spatial scales is the non-availability of species richness and distribution data for many taxa. Most of the existing global studies tend to focus on plants (Massante et al., 2019), birds (Jetz et al., 2014), mammals (Safi et al., 2011), and herpetofauna (Fritz & Rahbek, 2012) but have largely ignored arthropod taxa (Beck & McCain, 2020). Among arthropods, predatory soil-dwelling communities have been particularly neglected in macroecological studies (Finch, Blick & Schuldt, 2008), as they typically consist of many cryptic species occurring in low abundance making them difficult to detect. This is coupled with a lack of taxonomic expertise in identifying them to the specieslevel, as well as the presence of many undescribed species/lineages. In many ecosystems, predatory soil arthropods are likely to be the oldest lineages and play an important role in maintaining the ecosystem, but their diversity patterns remain poorly understood. Centipedes (class: Chilopoda) are one such group of soil arthropods, which represent one of the four main myriapod lineages, with a 420 million year (Ma) old fossil history that makes them some of the oldest living terrestrial predators (Edgecombe & Giribet, 2019). Among centipedes, the family Scolopendridae from the Western Ghats (WG) biodiversity hotspot in peninsular India is a well-characterized community, which offers a unique opportunity for conducting macroecological studies (Fig. 1).

**Figure 1.**
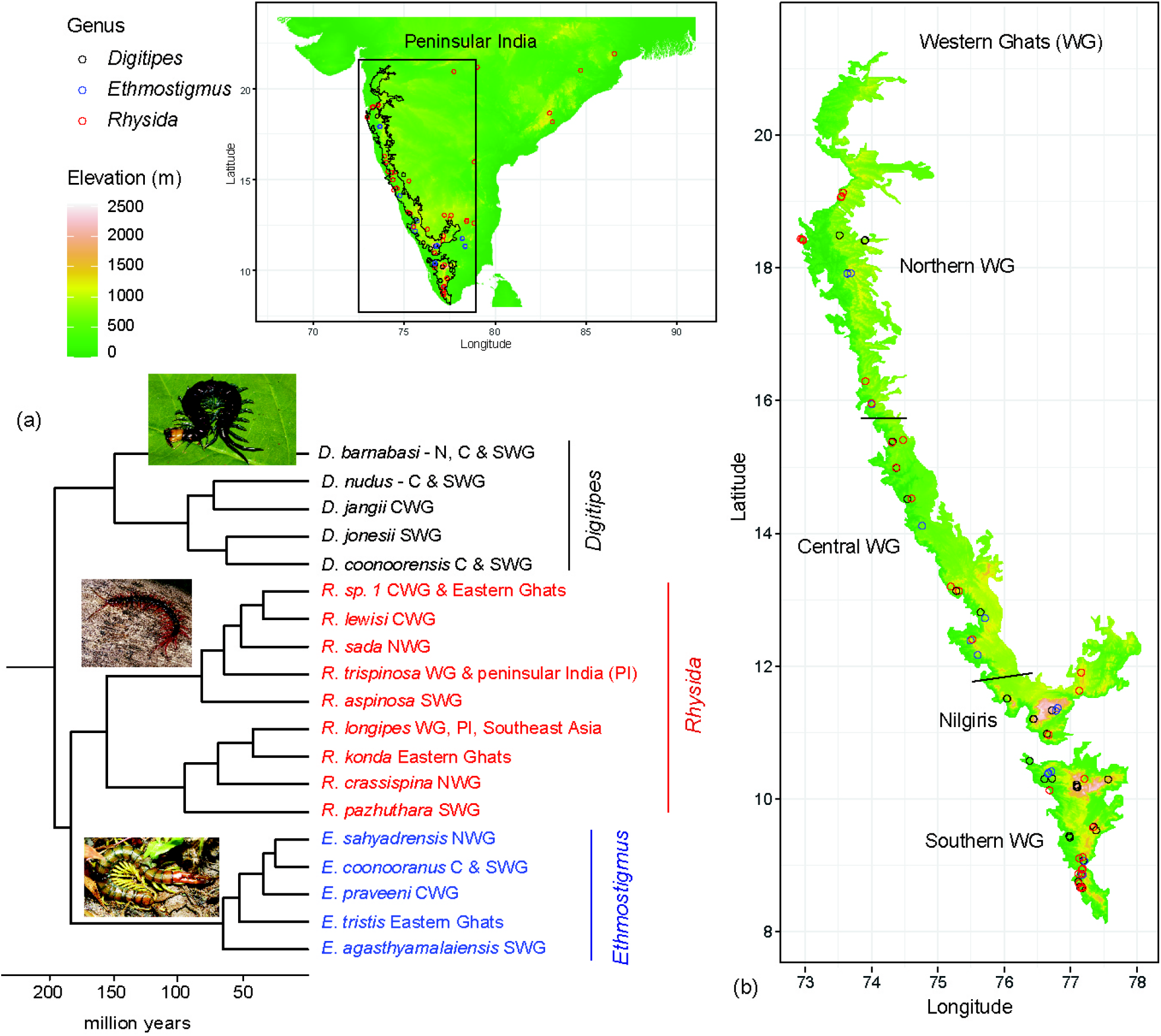
(a) The Bayesian phylogeny (b) A detailed map of the distribution records across peninsular India with a focus on scolopendrid centipede communities of the Western Ghats.

The scolopendrid community is one of the oldest (Late Cretaceous - 100 Ma) soil arthropod communities in the WG, and among the most diverse centipede groups in tropical Asian forests. The WG scolopendrid community has been revised extensively by using both morphology and molecular data, resulting in the discovery of many endemic species and radiations (Joshi & Edgecombe, 2018; Joshi, Karanth & Edgecombe, 2020). It is accompanied by detailed primary distribution data spanning large latitudinal (8°N - 20°N) and elevational gradients (100 - 2400 meters from mean sea level (msl)). It is noteworthy that molecular data were integrated with morphology and species ecology in these studies, as the latter often receive lesser attention in the current barcoding era (Padial et al., 2010). The scolopendrid centipedes of the WG vary in their endemicity patterns, ranging from narrow-range endemics to species with wide distributions across the WG or, exceptionally widespread distribution into SE Asia. In this study, we focus on a data-set representing one of the scolopendrid subfamilies, Otostigminae, which is useful for assessing hypotheses related to the patterns of diversity and endemism for predatory soil arthropods in tropical forests at a community level. Globally such datasets are rare, most of them consisting of island communities (Caribbean: Crews & Esposito, 2020; Azores: Borges & Hortal, 2009; Galapagos: Peck, 2006; Hawaii: Gillespie, 2002).

The Western Ghats (WG) is a 1600 km (8°N – 21°N) long mountain chain that runs along the west coast of peninsular India. The WG has been identified as a global biodiversity hotspot due to its high diversity and endemicity (Myers et al., 2000), and a recent biodiversity assessment reported that 30% of India’s biodiversity is found in the WG with a high proportion of endemic species (CEPF, 2016). This mountain chain has been divided into four phytogeographic subregions corresponding to northern WG (river Tapi to Goa), central WG (river Kali to Coorg), Nilgiris and the southern WG (Anamalai, Palani and Cardamom hills) (Subramanyam & Nayar, 1974) (Figure 1). The WG mountain chain has a prominent 30 km-wide break at 11°N, known as the Palghat Gap (Figure 1), which separates the Nilgiris from the southern WG and is thought to be an important biogeographic barrier (Joshi & Karanth, 2013; Robin et al., 2015; Vijayakumar et al., 2016).

Biogeographically, WG is a complex landscape as it was part of the Gondwanan supercontinent ca. 200 Ma ago and merged with Asia only recently, ca. 50 Ma. As a result, the WG harbours taxa with both Gondwanan and Asian affinities as well as many endemic radiations (Bossuyt & Milinkovitch, 2001; Gower et al., 2011; Joshi & Edgecombe, 2019; Joshi & Karanth, 2011; Surveswaran et al., 2020). In addition, peninsular India also experienced massive volcanic activity during the Cretaceous (ca. 65 Ma), which triggered mass extinctions. However, the southern WG remained climatically stable during this volcanic activity and is thought to have acted as a refugium for wet evergreen species (SWG refuge hypothesis - Joshi & Karanth, 2013 and references therein). The SWG refuge hypothesis predicts that older lineages will be present in the SWG and younger lineages will be present in the CWG and NWG. In addition to the geo-climatic processes, there is also a seasonality gradient along the WG, with northern latitudes showing greater temperature and precipitation seasonality than the southern latitudes, which has been shown to influence diversity patterns (Joshi & Karanth, 2013; Bose et al., 2019; Page & Shanker, 2020). However, an explicit examination of the diversity patterns using distributional, climatic and phylogenetic data remains to be explored for WG biota.

In this study, we aimed to understand how the diversity of the centipede community is structured in the WG. Based on distribution and biogeographic studies of plants and animals from this region (plants: Davidar et al., 2005, Page & Shaker, 2020; snails: Aravind et al., 2005; frogs: Daniels, 1992, Aravind & Gururaja, 2011), we expected a latitudinal diversity gradient (LDG) in which diversity increases from higher to lower latitudes in the WG. Additionally, the SWG refuge hypothesis provides expectations of ancient and high diversity in the southern WG and Nilgiris, as a result of climatic stability allowing more time for speciation, while central and northern WG taxa have relatively younger (<65 Ma) and fewer species. To explore these questions, we used species distribution models to map the spatial patterns of diversity and endemism instead of relying only on point locations from sampling surveys. We then used predictions from species distribution models to compare patterns of taxonomic diversity and endemism with phylogenetic diversity and endemism to identify areas with unique diversity. We asked the following specific questions –

1. How is centipede diversity distributed in the WG? Based on the LDG and SWG refuge hypotheses, we expect to see a decreasing gradient in diversity from southern to northern WG.
2. Are there hotspots within the hotspot? Since species diversity may not be uniformly distributed given the climatic and topographic heterogeneity in the WG, we examined if there are areas represented by disproportionately high diversity and endemism within the WG biodiversity hotspot.
3. What are the patterns of species turnover across the WG? We assessed if taxonomic and phylogenetic composition are unique to each of the biogeographic subregions in the WG.

## Methods

### Species distribution models

Primary location data (n = 100) for 19 species in the three genera of the centipede family Scolopendridae (Subfamily Otostigminae): *Digitipes* Attems, 1930, *Ethmostigmus* Pocock, 1898, and *Rhysida* Wood, 1862 were obtained by systematic sampling across the Western Ghats (WG) from 2008 to 2010, spanning its latitudinal and elevational gradients (Joshi et al., 2020; Joshi & Edgecombe, 2013, 2018; Joshi & Karanth, 2012) (see Appendix S1 in Supporting Information). These data were supplemented with opportunistic sampling which continued to 2018. These locations spanned the extent of peninsular India, with a focus on the wet forests of the WG, a global biodiversity hotspot (Fig. 1B). Since it is challenging to identify centipede species in the field, specimens associated with the primary location data were collected and identified in the lab based on microscopic examination of morphological characters. Species identity was also assessed through molecular phylogenetic and species delimitation analyses (Joshi et al., 2020; Joshi & Edgecombe, 2013, 2018; Joshi & Karanth, 2012). State forest department permits were obtained to collect centipedes in protected areas and specimens were preserved in 70% ethanol. A few secondary locations (n = 10) were obtained from published sources, where we were certain about the species identification based on morphological characters described in the source literature (Jangi & Dass, 1984).

It is difficult to determine true absence for a group such as centipedes due to their low abundance, morphologically cryptic nature and lack of systematic information on their distribution in less explored areas such as the WG. Therefore, we chose to model species distributions using Maxent version 3.4.1 (Phillips et al., 2006), which uses a presence-only approach to predict species distributions that has been shown to perform well across species, regions (Elith et al., 2006; Phillips & Dudík, 2008) and a range of sample sizes (Hernandez et al., 2006; Wisz et al., 2008). Maxent compares the environment at presence locations against background locations drawn from the model extent to arrive at a model of relative suitability for a species based on the underlying environmental variables (Elith et al., 2011; Merow et al., 2013). We ran Maxent models for each species for the model extent of peninsular India (8° - 24°N, 68° - 91°E) at 30 arc second resolution (0.0083 × 0.0083 degree resolution, 0.93 × 0.93 km at equator). We used presence locations from each species, 10,000 background locations derived from a sampling bias model, predictor variables consisting of environmental layers from the WorldClim database (Fick & Hijmans, 2017) and a soil type layer (ATREE Spatial Archive, 2020) to run models using default parameters. We used mean and standard error of Area Under the Receiver Operator Curve (AUC) (Fielding & Bell, 1997) and True Skill Statistic (TSS) (Allouche et al., 2006) obtained from five-fold cross-validation, and a jack-knifing test for small sample sizes (Pearson, 2007) to evaluate model predictions (details of the Maxent modeling approach and evaluation are provided in Appendix S2).

Spatial data were processed using the packages ‘rgeos’ (Bivand & Rundel, 2017), ‘raster’ (Hijmans, 2020) and ‘sp’ (Bivand, Pebesma & Gomez-Rubio, 2013; Pebesma & Bivand, 2005), and Maxent models were run using the package ‘dismo’ (Hijmans et al., 2017) in R 3.6.1 (R Core Team, 2019).

### Diversity and endemism measures

The diversity and endemism measures derived from these predicted distributions at the scale of peninsular India were cropped for the WG for further analysis, since this biodiversity hotspot is the focus of our study where systematic sampling was undertaken. We aggregated continuous Maxent predictions of relative habitat suitability to obtain maps at a scale of 0.83 × 0.83 degrees (93 × 93 km at equator). We assigned the maximum value of habitat suitability among the underlying cells to the larger aggregated cell and applied a threshold of maximum sum of sensitivity and specificity (Liu et al., 2005) to convert it into a presence-absence map for each species.

We used these binary maps to calculate taxonomic diversity (TD) and weighted endemism (WE) along with their phylogenetically informative counterparts – phylogenetic diversity (PD) and phylogenetic endemism (PE) for each cell in the model extent. As compared to taxonomic indices of diversity and endemism, the phylogenetic indices provide additional information on evolutionary relationships between species present within a community, helping us distinguish between closely and distantly related species. The diversity indices enable us to test the latitudinal diversity gradient hypothesis in the WG, where we expect to observe increasing diversity with decreasing latitude. Combined with the endemism indices, they help us test predictions from the southern WG refuge hypothesis where we would expect to see high diversity and endemism within this sub-region of the WG. Additionally, both the indices of endemism allow us to identify hotspots consisting of range-restricted species within the WG, where PE additionally identifies evolutionarily unique and range-restricted species.

TD is calculated by stacking species distributions and summing species presences for each cell, while WE is calculated by scaling species presence with its range size (number of cells in the predicted map where a species is present) and summing this across species found in a cell (Crisp et al., 2001). A species time-tree for the subfamily Otostigminae based on a combined dataset of mitochondrial and nuclear markers (Joshi et al., 2020) was used to calculate PD and PE. PD is calculated by summing up branch lengths of all species present in a cell derived from the phylogenetic tree (Faith, 1992) and we present it as the proportion of total tree length (range: 0 - 1). PE additionally scales the branch length with range size prior to summation across species in a cell (Rosauer et al., 2009) and we also present it as a proportion of total tree length (range: 0 – 1).

To compare and identify differences in the relative distribution of evolutionary ages among lineages present within different communities, we used relative phylogenetic diversity (RPD) and relative phylogenetic endemism (RPE). These indices compare the observed values of PD and PE with those obtained from a phylogenetic tree with equal branch lengths, which allows us to understand if there is an over-representation of evolutionarily old or young lineages within a community. This information can provide insights into the biogeographic history or ecological processes operating in a region, for example, it can help us distinguish between centres of neo- and paleo-endemism (Mishler et al., 2014).

RPD and RPE were calculated as the ratio of PD and PE derived from the actual phylogenetic tree over the same indices derived from a phylogenetic tree where species relationships remain the same but lineages have equal branch lengths. An RPD/RPE value of 1 indicates that the lineages present in a cell have equal branch lengths, RPD/RPE values larger than 1 indicate regions harbouring species with longer branch lengths in the phylogenetic tree, and values smaller than 1 indicate regions harbouring species with shorter branch lengths in the phylogenetic tree (Mishler et al., 2014).

We generated null distributions of phylogenetic diversity and endemism measures for comparison with observed values by randomizing tip labels on the phylogenetic tree while keeping the taxonomic diversity of each cell and range size of each species constant (Mishler et al., 2014). The position of observed phylogenetic diversity and endemism values was described as percentiles with respect to the resulting null distribution. Further examination of the endemism patterns using the CANAPE pipeline (Mishler et al., 2014) is reported in Appendix S3. The diversity and endemism indices, and their corresponding null distributions were calculated using Biodiverse 3.1 (Laffan et al., 2010).

We used measures of beta-diversity to assess if taxonomic and phylogenetic composition varies between the different biogeographic sub-regions recognized along the WG. To identify patterns of taxonomic beta diversity, we used the Simpson dissimilarity index, which describes the variation in species composition due to species turnover alone (Baselga, 2010). We also calculated its phylogenetic counterpart in the form of PhyloSor_Turn_, which measures the loss of branch lengths between communities not explained by differences in phylogenetic diversity (Leprieur et al., 2012). The taxonomic and phylogenetic indices of turnover were used for cluster analysis using the UPGMA algorithm (Michener & Sokal, 1957) to group sub-regions in the WG (*k* = 4) based on patterns of species composition. Beta diversity and phylogenetic beta diversity indices were calculated using the ‘betapart’ package (Baselga, 2010) in R 3.6.1 (R Core Team, 2019).

## Results

### Maxent predictions for the scolopendrid community

Most (17 out of 19) species distribution models had AUC values above 0.8, and all of them had TSS values above 0, indicating that they performed better than random classification. The Maxent model for *Rhysida longipes* had an average AUC value of 0.3771, but an acceptable TSS value of 0.2758 and its jackknifing test statistic was significantly different from a random model (Table 1). Among species with small sample sizes, *Rhysida* sp. 1 had acceptable AUC and TSS values, though the jackknifing test could not differentiate model predictions from random assignment (Table 1). Therefore, we used available presence locations instead of model predictions for diversity and endemism calculations for this species. Maxent results showed that variables related to precipitation and temperature seasonality and maximum temperature were important in predicting the distributions of centipede species (see Appendix S2).

**Table 1.**
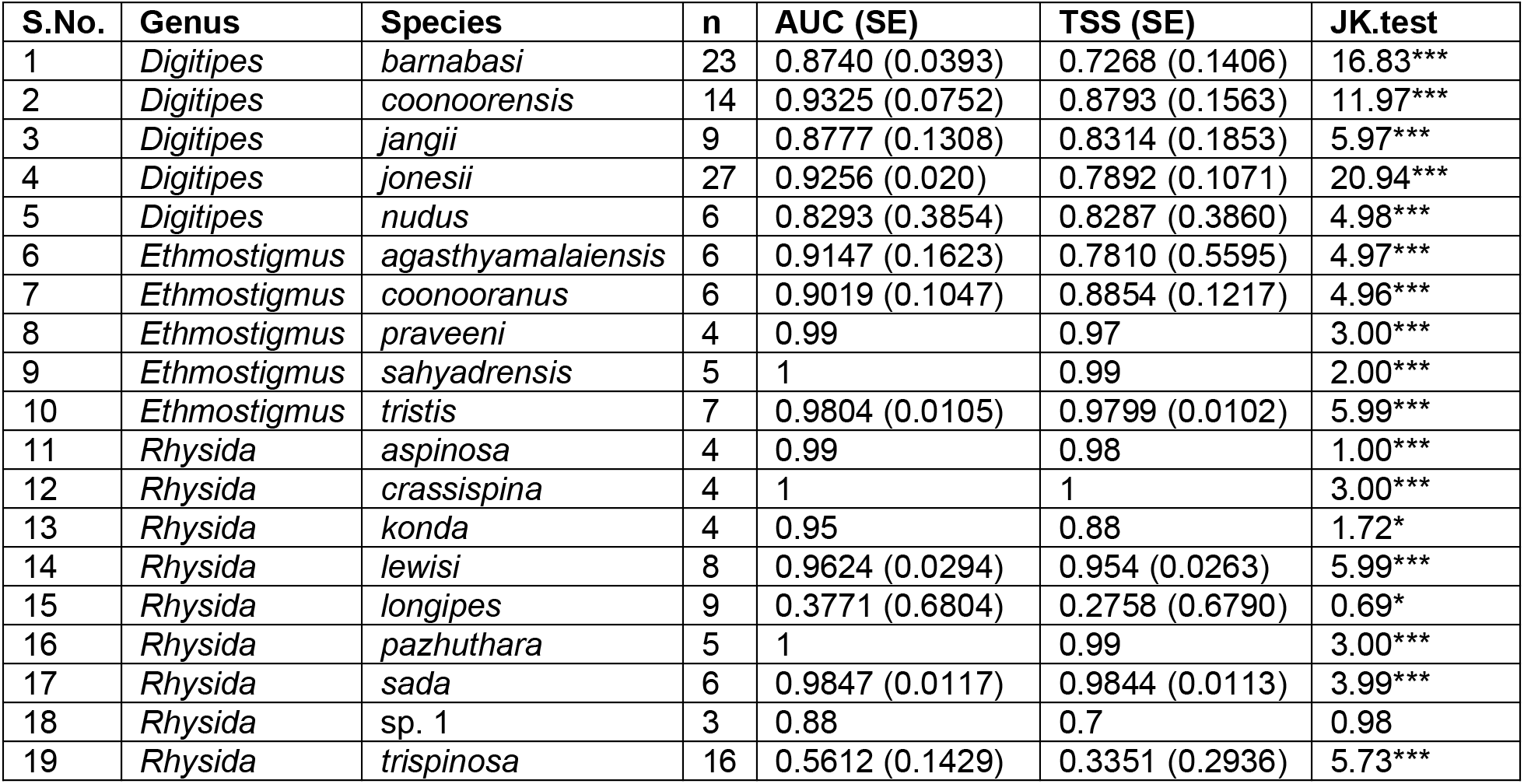
Model evaluation of Maxent species distribution models run at 0.0083° × 0.0083° resolution for peninsular India. AUC – Area Under the Receiver Operator Curve, TSS – True Skill Statistic, JK.test – Test statistic for jack-knifing test for small sample sizes. For species with greater than five presence locations, mean and standard error (within parentheses) of evaluation metrics are presented from five-fold cross-validation. \**p*<0.05, \*\**p*<0.01, \*\*\**p*<0.001

### Patterns of diversity and endemism

Patterns of taxonomic and phylogenetic diversity were found to be broadly concordant for all four subregions, namely southern WG, Nilgiris, central and northern WG (Figs 2a, 2b). The differences between these measures were mainly in the relative magnitude of diversity values. The southern WG had the highest values of phylogenetic diversity indices and also higher than expected values of phylogenetic diversity (Fig. 2b) and relative phylogenetic diversity (Fig. 2c) as compared to the null distribution. Higher than expected values of relative phylogenetic diversity indicate the presence of a greater proportion of lineages with long-branch lengths. Nilgiris can be ranked next in terms of phylogenetic diversity values (Fig. 2b). Central WG followed next in the magnitude of phylogenetic diversity (Fig 2b), but had lower than expected values of relative phylogenetic diversity (Fig. 2c) indicating the prevalence of species with short branch lengths. The northern WG had the lowest phylogenetic diversity (Fig. 2b) but high values of relative phylogenetic diversity (Fig. 2c).

**Figure 2.**
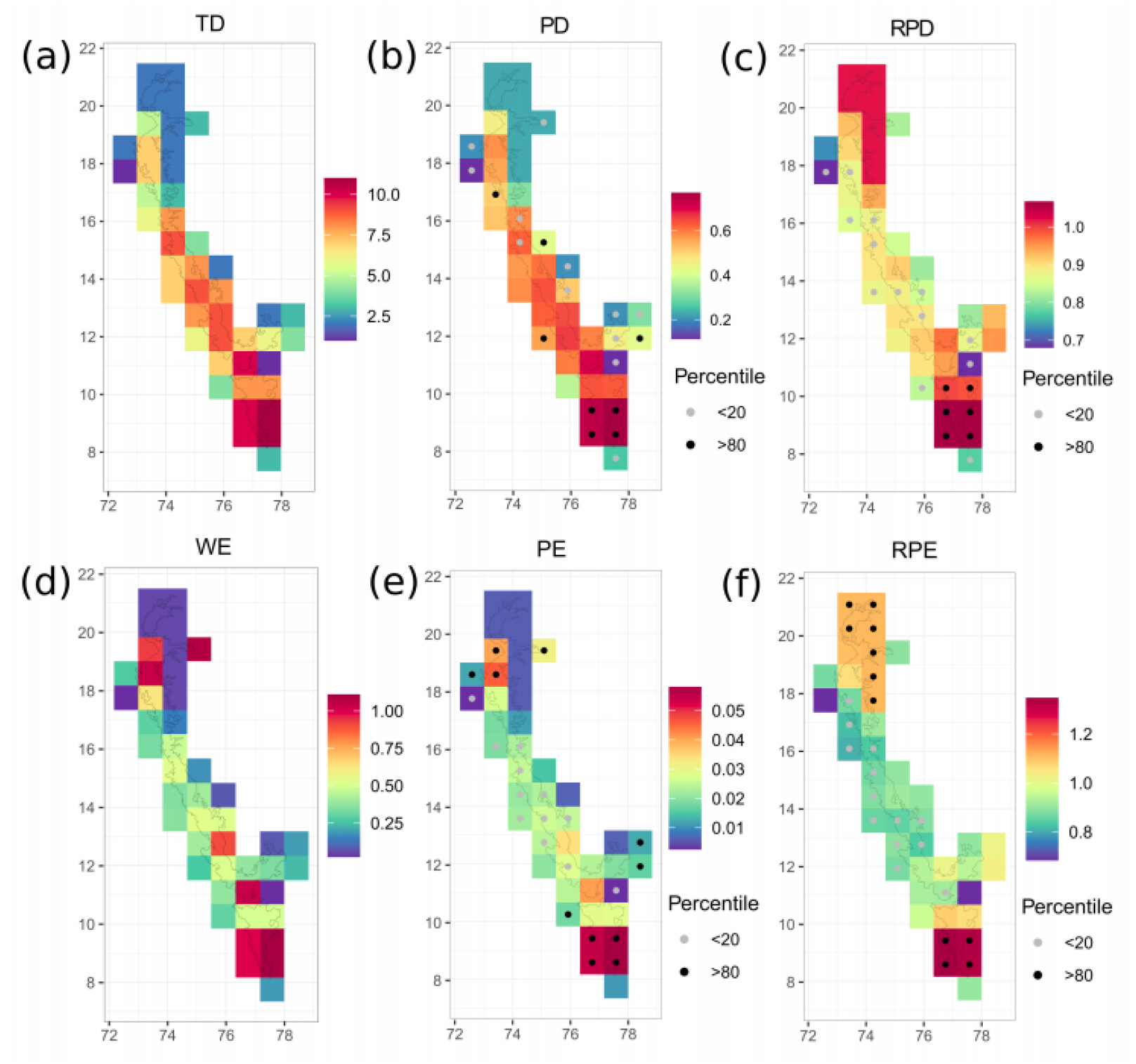
Maps of diversity and endemism indices for scolopendrid centipedes in the Western Ghats, India calculated at a resolution of 0.83° × 0.83° grid cells (a) taxonomic diversity presented as absolute counts of species in each grid cell, (b) phylogenetic diversity presented as proportion tree length where large values correspond to greater phylogenetic diversity, (c) relative phylogenetic diversity where values greater/lesser than 1 indicate greater relative proportion of older/younger lineages, (d) weighted endemism (taxonomic) where larger values correspond to greater species endemism, (e) phylogenetic endemism presented as proportion tree length where large values correspond to greater phylogenetic endemism and (f) relative phylogenetic endemism where values greater/lesser than 1 indicate greater relative proportion of older/younger endemic lineages. For the phylogenetic indices, filled circles within cells indicate the percentile rank of the observed index within a null distribution of index values obtained by shuffling tip labels on the phylogenetic tree.

When taxonomic and phylogenetic diversity were informed by species geographic ranges, regions in the southern WG converged in having both high and higher than expected values of weighted and phylogenetic endemism (Figs 2d, 2e). Southern WG was also associated with higher than expected values of relative phylogenetic endemism (Fig. 2f), indicating that the species found here are characterized by small range sizes and a higher proportion of lineages with long branch lengths. The central WG had low to intermediate values of phylogenetic endemism and lower than expected values of relative phylogenetic endemism, indicating a higher proportion of species with relatively wide distributions and short branch lengths. Most areas within the northern WG, including regions which showed variation in phylogenetic endemism, emerged as having higher than expected values of relative phylogenetic endemism (Fig. 2f).

The patterns of diversity and endemism calculated using binary maps at the larger spatial scale broadly correspond with values derived from habitat suitability predictions in Maxent at the native resolution without applying a threshold (see Appendix S4), which is thought to more closely estimate these indices (Calabrese et al., 2014).

### Patterns of species composition

Simpson dissimilarity and PhyloSor_Trun_, which describe the turnover component of beta diversity, grouped the southern WG and Nilgiris separately from central and northern WG. Some of the cells in the eastern edge of the central and northern WG were also a part of this group, which might be due to the predicted distribution patterns of some peninsular Indian species (such as *E. tristis* and *R. trispinosa*) extending into the WG (Figs 3a, 3b).

**Figure 3.**
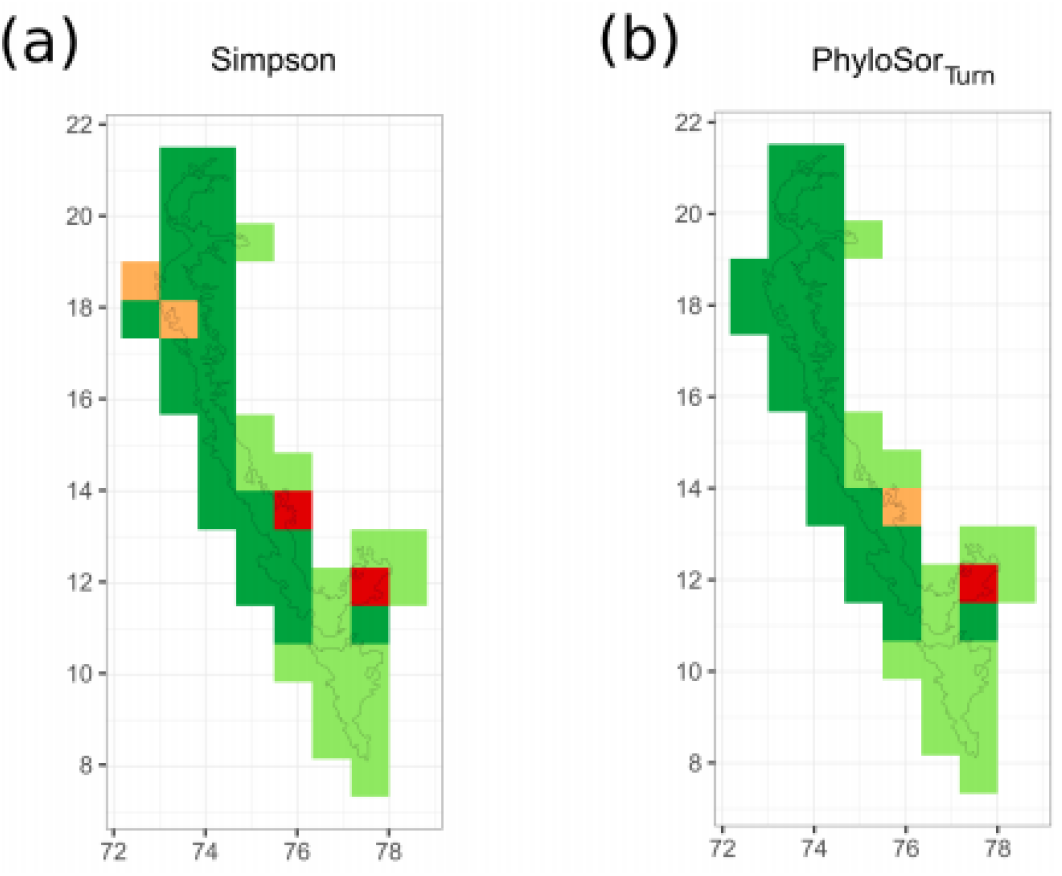
Spatial clusters recovered from UPGMA trees (*k* = 4) of pairwise distances related to (a) Simpson dissimilarity (species turnover) and (b) phylogenetic turnover (PhyloSor_Turn_) in the Western Ghats, India in 0.83° × 0.83° grid cells. Colours on the map represent membership of each grid cell to each of the four clusters recovered.

## Discussion

This is one of the first studies that encompasses the entire stretch of the WG to systematically evaluate the patterns of diversity, endemism and composition using primary distribution data to model species distributions along with a robust, dated species-level phylogeny. Measures of taxonomic and phylogenetic diversity show support for predictions from the latitudinal diversity gradient (LDG) hypothesis, where we expect centipede diversity to decrease from lower to higher latitudes in the WG. This finding also supports expectations from the southern refuge hypothesis, where we expect the southern WG to have higher taxonomic and phylogenetic diversity than the other WG sub-regions. We also found that weighted and phylogenetic endemism broadly converged in the WG, where regions of high endemism were detected in the southern WG, and in plateaus with stunted forests in the northern WG.

Taxonomic diversity and all the indices of phylogenetic diversity were higher in the southern WG as compared to other regions. The southern WG is followed by Nilgiris and central WG in the magnitude of phylogenetic diversity and both these regions have lower values of relative phylogenetic diversity as compared to southern WG. Northern WG has the lowest phylogenetic diversity but shows higher values of relative phylogenetic diversity suggesting presence of unique lineages.

The decline in diversity with increasing latitude is a well-known pattern across taxa and regions (Hillebrand, 2004; ants: Economo et al., 2018; angiosperms: Kerkhoff, Moriarty & Weiser, 2014; mammals: Rolland et al., 2014). Various explanations have been proposed to explain LDG, where higher species richness at lower latitudes has been attributed to greater time available for speciation, higher speciation rates or lower extinction rates (Mittelbach et al., 2007). The causal drivers of variation in speciation and extinction rates can be related to geographic area, productivity, time, climatic stability, temperature and biological interactions (Fine, 2015). In the case of the WG, we believe geological events and climatic gradients might explain the observed diversity patterns. The southern WG may have acted as a refuge during Cretaceous volcanism (ca 65 Ma), which is thought to be associated with widespread extinctions of plant and animal species in the northern WG (Joshi & Karanth, 2013 and references therein). This, along with climatic stability over long periods of time as inferred from past vegetation patterns (Prasad et al., 2009; Divya and Karanth), could have led to the persistence and diversification of ancient centipede lineages in the southern WG (Divya et al., 2020). On the other hand, higher latitudes of the WG are observed to have increasing seasonality (Page & Shanker, 2020), which is associated with the Miocene emergence of the Indian monsoon (Gunnell, 1997) and has been implicated in the decline in taxonomic (Page & Shanker, 2020) and phylogenetic diversity in plants (Bose et al., 2019).

### Southern Western Ghats – hotspot within a hotspot

We show that the centipede communities of the southern WG not only have high taxonomic diversity but also high phylogenetic diversity. A similar pattern was reported for one of the centipede genera *Digitipes* (Joshi & Karanth, 2013), and the current study shows that this pattern holds true even after adding an additional 14 species from two other genera, highlighting the importance of southern WG with respect to centipede diversity. Examination of relative phylogenetic diversity and endemism patterns allowed us to identify ancient, evolutionarily and geographically unique lineages in the southern WG. High taxonomic diversity in the southern WG has been documented in plants (Page & Shanker, 2020) and animals (Aravind & Gururaja, 2011; Aravind et al., 2005; Daniels, 1992). However, an understanding of diversity patterns from a phylogenetic perspective remains limited and needs to be explored further. Given the complex geological history of the WG, it is also likely that there were multiple and/or different refugia within the southern WG over different time periods (Joshi & Karanth, 2013; Bose et al., 2019). The use of a dated phylogeny in calculating diversity and endemism measures allowed us to identify ancient endemic centipede lineages in the southern WG, which points to its role as a refugium during Cretaceous volcanic activity. Additionally, lower climatic seasonality in the southern WG might have contributed to higher diversity in this subregion, which needs to be investigated in greater detail in the future.

There are mixed signals from studies from other regions and taxa, which have compared the relative importance of current and past climate as well as historic geography in driving patterns of diversity and endemism. For example, in millipedes of south-west Australia, landscape evolution is thought to be an important driver of richness and endemism patterns in some orders, whereas overall species turnover is related to contemporary rainfall patterns (Moir et al., 2009). In orchid bees of Amazonian rainforests, climatic seasonality is associated with reduced phylogenetic diversity at higher latitudes, while past geological events play a more important role in shaping bee diversity at lower latitudes (Abrahamczyk et al., 2014). Our results from soil arthropods highlight the need for macroecological analyses on a diverse range of taxa to understand the relative importance of geological processes and climatic variables in governing diversity and endemism patterns in the WG

### Less diverse but unique northern Western Ghats

While the northern WG had the lowest phylogenetic diversity it still had fairly higher values of relative phylogenetic endemism, though not significant indicating the presence of relatively older endemic lineages along with recent endemic lineages within its communities. We also noted a relative increase in phylogenetic endemism from central to some cells in the northern WG, which is related to the presence of narrow endemic species such as *R. crassispina, R. sada*, and *E. sahyadrensis*. These species are found in unique habitats consisting of stunted evergreen forests and adjoining grasslands on lateritic plateaus in the northern WG. These plateaus are geologically unique and are made up of basalt from Deccan Trap lava flows, which has weathered into lateritic rock that has further undergone various levels of erosion (Watve, 2013). They show a diversity of unique seasonal microhabitats (Thorpe et al., 2018) and have a distinct vegetation consisting of several endemic herbaceous species that show adaptations to surviving in poor soil conditions (Joshi & Janarthanam, 2004; Lekhak & Yadav, 2012).

Unfortunately, the network of protected areas in the northern WG is not as extensive as in the southern WG, though they consist of areas identified to be of high conservation value (Das et al., 2006; Watve, 2013). The protected areas in the northern WG are small in size and consist of fragmented forests with high anthropogenic disturbance located in the vicinity of urban centers (Gadgil et al., 2011; Thorpe & Watve, 2015). Apart from centipedes, there have been records of other range-restricted species on these plateaus across different taxa (snails: Aravind et al., 2005; plants: Lekhak & Yadav, 2012; Shigwan et al., 2020; amphibians: Katwate et al., 2013). The present study highlights the urgent need to systematically understand the evolutionary uniqueness of species found in the plateau across different taxa and identify key areas of conservation importance.

### Role of the Palghat Gap in structuring centipede communities

The Palghat Gap, which is 30 km wide valley interrupting the otherwise contiguous Western Ghats mountain chain, has been identified as an important barrier for dispersal (Joshi & Karanth, 2013; Robin et al., 2015; Subramanyam & Nayar, 1974; Vidya et al., 2005). Interestingly, taxonomic and phylogenetic turnover in centipedes revealed two major clusters, which were largely restricted to either the north or south of the Palghat Gap, suggesting species replacement across this valley. This highlights the role of biogeographic barriers in shaping diversity and patterns of community composition. Species from the WG which spanned the Palghat Gap based on observed occurrence locations, include *Digitipes barnabasi, D. coonoorensis, D. jonesii, D. nudus, Ethmostigmus coonooranus, Rhysida longipes* and *R. trispinosa*, and they also spanned this potential barrier based on their predicted distribution.

There is increasing evidence that biogeographic barriers in addition to the climatic barriers can shape community dynamics across tropical areas. In the tropical Andes, river valleys and elevation have been shown to drive distribution and phylogenetic breaks in endemic bird taxa. These barriers are found to encompass areas with high richness of narrowly distributed species (Hazzi et al., 2018). Major rivers also demarcate bioregions which explain distribution patterns of anurans in Amazonia, which is followed in importance by the climatic and topographic variation (Godinho & da Silva, 2018). In the Australian monsoon tropics, biogeographic barriers have shaped the distribution patterns for plants and several animal taxa (Edwards et al., 2017). Our results recommend the simultaneous assessment of geo-climatic factors while examining patterns of diversity and endemism in the Western Ghats, given its complex geological past and the contemporary gradient in temperature and precipitation seasonality.

### Directions for centipede distributions within and outside the Western Ghats

Among the centipede genera studied here, in India, *Digitipes* is restricted to the wet forests of the Western Ghats, while *Ethmostigmus* lineages have dispersed to wet forests in the Eastern Ghats in the past (Joshi & Edgecombe, 2019) and *Rhysida* is widely distributed across different habitat types throughout peninsular India. Our species distribution models predict ranges of *E. tristis, R. konda* (both of which are Eastern Ghats endemics) and the more widespread *R. trispinosa*, which extend into the eastern boundary of the Western Ghats. The presence of these species in their range extremes along with species having more extensive distributions across this mountain range may lead to unique centipede communities in the eastern edge of the Western Ghats.

The niche modelling predictions on relative habitat suitability can serve as a useful guide for sampling effort to assess population-level patterns and processes in centipedes of the Western Ghats. The predicted distributions of many species extend northwards from their observed presence locations, which needs to be investigated to confirm their range boundaries (see Appendix S2). These include distributions of species from northern and central Western Ghats such as *D. coonoorensis, D. nudus, E. conooranus, E. praveeni, R. lewisi* and *E. sahyadrensis*, which are predicted to have distributions extending further north from currently reported occurrences. The models developed here can also be used to predict potential distributions in other under-sampled areas such as wet forests of the Eastern Ghats, and north-east India.

To conclude, we demonstrated the use of primary distribution data in conjunction with species distribution modelling and detailed species-level phylogeny to understand diversity gradients in tropical wet forests in an ancient soil arthropod group (centipedes). Our work indicates that a combination of current and past climatic factors as well as geography has shaped the diversity and distributions of centipedes in the WG. Future studies need to be carried out at the level of the soil arthropod community and also other lesser studied taxa in the WG to understand diversity dynamics in these diverse tropical forests. This approach involving both ecological and evolutionary factors also promises to be useful in identifying areas of endemism across taxa within the biodiversity hotspot, which is an important exercise while identifying areas for conservation importance.

## iv. Acknowledgements

D. K. Bharti was supported during this study by a start-up grant to JJ from CSIR-Centre for Cellular and Molecular Biology, Uppal Road, Hyderabad, India. We would like to thank Dr Rohit Naniwadekar, Dr Navendu Page and Abhishek Gopal for insightful discussions and comments on the paper.

## v. Biosketch

D. K. Bharti is a postdoctoral researcher interested in processes shaping patterns of species distribution and genetic diversity, especially in poorly studied tropical terrestrial and marine invertebrates.

Jahnavi Joshi is an evolutionary ecologist studying arthropod systematics and examining the evolutionary and ecological processes that shape arthropod diversity in Asian tropical forests.

Praveen Karanth is a phylogeneticist interested in using phylogenies to address questions in the areas of ecology, evolution and behaviour.

Gregory D. Edgecombe is a systematist investigating the higher level phylogenetics of arthropods and the taxonomy and evolution of centipedes.

## Data Accessibility Statement

The location data used in this study are provided in Appendix S1 of Supporting Information. The environmental layers used for species distribution modeling are available for download from https://www.worldclim.org/data/bioclim.html. R scripts used for building species distribution models, generating input files for Biodiverse 3.1, calculating indices, and generating plots are available at https://github.com/bhartidk/centipede_diversity_endemism.

---

**Appendix S1**: A list of geographic locations for 19 scolopendrid centipede species used for the species distribution models.

**Appendix S2:** Description of Maxent species distribution modeling approach and results.

**Appendix S3:** Identifying hotspots of phylogenetic endemism using the Categorical Analysis of Palaeo and Neo Endemism (CANAPE).

**Appendix S4:** Diversity and endemism measures calculated using continuous predictions of habitat suitability from Maxent species distribution models.

