## Supporting Information Appendix S1 for "Spatial patterns of phylogenetic diversity and endemism in the Western Ghats, India: a case study using ancient predatory arthropods"

**Appendix 1.** A list of geographic locations for 19 Scolopendrid centipede species used for the species distribution models.

| **Genus** | **Species** | **Location** | **Latitude** | **Longitude** | **Reference** |
| --- | --- | --- | --- | --- | --- |
| *Digitipes* | *barnabasi* | Bopdev ghat, Pune district, Maharashtra, India | 18.4189 | 73.9068 | Joshi & Karanth 2012 |
| *Digitipes* | *barnabasi* | Castle-rock, Uttara Kannada district, Karnataka, India | 15.3698 | 74.3063 | Joshi & Karanth 2011 |
| *Digitipes* | *barnabasi* | Kumta, Uttara Kannada district, Karnataka, India | 14.5257 | 74.6025 | Joshi & Karanth 2012 |
| *Digitipes* | *barnabasi* | Dudhsagar Water Falls, Goa-Karnataka border, South Goa district, Goa, India | 15.3868 | 74.3192 | Joshi & Karanth 2012 |
| *Digitipes* | *barnabasi* | Pune district, Maharashtra, India | 18.4103 | 73.8985 | Joshi & Edgecombe 2018 |
| *Digitipes* | *barnabasi* | Pune district, Maharashtra, India | 18.4145 | 73.9036 | Joshi & Karanth 2012 |
| *Digitipes* | *barnabasi* | Bombay point, Mahabaleshwar Reserve Forest, Satara district, Maharashtra, India | 17.9157 | 73.6363 | Joshi & Karanth 2012 |
| *Digitipes* | *barnabasi* | Amboli, Sindhudurg district, Maharashtra, India | 15.9594 | 73.9997 | Joshi & Karanth 2012 |
| *Digitipes* | *barnabasi* | Amboli, Sindhudurg district, Maharashtra, India | 15.9617 | 74.0021 | Joshi & Karanth 2012 |
| *Digitipes* | *barnabasi* | Talacauvery, Kodagu district, Karnataka, India | 12.4055 | 75.5199 | Joshi & Karanth 2012 |
| *Digitipes* | *barnabasi* | Bhimashankar, Pune district, Maharashtra, India | 19.0633 | 73.5396 | Joshi & Karanth 2012 |
| *Digitipes* | *barnabasi* | Bhimashankar, Pune district, Maharashtra, India | 19.0777 | 73.5505 | Joshi & Karanth 2012 |
| *Digitipes* | *barnabasi* | Bhimashankar, Pune district, Maharashtra, India | 19.1319 | 73.5708 | Joshi & Karanth 2012 |
| *Digitipes* | *barnabasi* | Bhimashankar, Pune district, Maharashtra, India | 19.1321 | 73.5706 | Joshi & Karanth 2012 |
| *Digitipes* | *barnabasi* | Bhimashankar, Pune district, Maharashtra, India | 19.1323 | 73.5701 | Joshi & Edgecombe 2018 |
| *Digitipes* | *barnabasi* | Devmali, Koraput district, Odisha, India | 18.6600 | 82.9900 | Joshi & Edgecombe 2018 |
| *Digitipes* | *barnabasi* | Sakaleshapur, Hassan district, Karnataka, India | 12.8133 | 75.6428 | Joshi, Karanth & Edgecombe, 2020 |
| *Digitipes* | *barnabasi* | Mulshi, Pune district, Maharashtra, India | 18.4916 | 73.5173 | Jangi & Dass, 1984 |
| *Digitipes* | *barnabasi* | Silent Valley National Park, Palakkad district, Kerala, India | 11.2089 | 76.4411 | Joshi & Karanth 2011 |
| *Digitipes* | *barnabasi* | Parakkadavu, Shendurney Wildlife Sanctuary, Kollam district, Kerala, India | 8.8619 | 77.1795 | Joshi & Karanth 2012 |
| *Digitipes* | *barnabasi* | Kurinjal, Kudremukh National Park, Chikkamagaluru district, Karnataka, India | 13.2008 | 75.1925 | Joshi & Karanth 2012 |
| *Digitipes* | *barnabasi* | Pasikada, Aachankovil Reserve Forest, Kollam district, Kerala, India | 9.0717 | 77.1916 | Joshi & Karanth 2012 |
| *Digitipes* | *barnabasi* | Kodagu district, Karnataka, India | 12.3889 | 75.4900 | Joshi, Karanth & Edgecombe, 2020 |
| *Digitipes* | *barnabasi* | Periyar Tiger Reserve, Idukki district, Kerala, India | 9.5755 | 77.3362 | Joshi & Karanth 2012 |
| *Digitipes* | *barnabasi* | Vellimalai, Periyar Tiger Reserve, Idukki district, Kerala, India | 9.5840 | 77.3490 | Joshi & Karanth 2012 |
| *Digitipes* | *coonoorensis* | Silent Valley National Park, Palakkad district, Kerala, India | 11.2022 | 76.4392 | Joshi & Karanth 2011 |
| *Digitipes* | *coonoorensis* | Silent Valley National Park, Palakkad district, Kerala, India | 11.2037 | 76.4387 | Joshi & Karanth 2011 |
| *Digitipes* | *coonoorensis* | Silent Valley National Park, Palakkad district, Kerala, India | 11.2089 | 76.4411 | Joshi & Karanth 2011 |
| *Digitipes* | *coonoorensis* | Siruvani Reserve Forest, Palakkad district, Kerala, India | 10.9820 | 76.6413 | Joshi & Karanth 2012 |
| *Digitipes* | *coonoorensis* | Waynad, Waynad district, Kerala, India | 11.5139 | 76.0434 | Joshi, Karanth & Edgecombe, 2020 |
| *Digitipes* | *coonoorensis* | Coonoor, Nilgiris district,Tamil Nadu, India | 11.3703 | 76.8041 | Jangi & Dass, 1984 |
| *Digitipes* | *coonoorensis* | Katteri, Ooty, Nilgiris district, Tamil Nadu, India | 11.3340 | 76.7172 | Jangi & Dass, 1984 |
| *Digitipes* | *coonoorensis* | Pambadum Shola National Park, Idukki district, Kerala, India | 10.1756 | 77.1062 | Joshi & Karanth 2012 |
| *Digitipes* | *coonoorensis* | Eravikulam National Park, Idukki district, Kerala, India | 10.1835 | 77.0897 | Joshi & Karanth 2012 |
| *Digitipes* | *coonoorensis* | Eravikulam National Park, Idukki district, Kerala, India | 10.1913 | 77.0974 | Joshi & Karanth 2012 |
| *Digitipes* | *coonoorensis* | Eravikulam National Park, Idukki district, Kerala, India | 10.2149 | 77.0944 | Joshi & Karanth 2012 |
| *Digitipes* | *coonoorensis* | Eravikulam National Park, Idukki district, Kerala, India | 10.2187 | 77.0894 | Joshi & Karanth 2012 |
| *Digitipes* | *coonoorensis* | Pandimatte, Shendurney Wildlife Sanctuary, Kollam district, Kerala, India | 8.8728 | 77.1634 | Joshi & Karanth 2012 |
| *Digitipes* | *coonoorensis* | Mesapulimalai, Idukki district, Kerala, India | 10.1756 | 77.1062 | Joshi & Edgecombe 2018 |
| *Digitipes* | *coonoorensis* | Palani, Dindigul district, Tamil Nadu, India | 10.2980 | 77.5640 | Jangi & Dass, 1984 |
| *Digitipes* | *coonoorensis* | Tadoli, Kudremukh National Park, Chikkamagaluru district, Karnataka, India | 13.1339 | 75.2782 | Joshi & Karanth, 2012 |
| *Digitipes* | *jangii* | Kali Tiger Reserve, Karwar district, Karnataka, India | 15.2725 | 74.9603 | Joshi & Karanth 2012 |
| *Digitipes* | *jangii* | Kali Tiger Reserve, Karwar district, Karnataka, India | 14.9878 | 74.3748 | Joshi & Karanth 2012 |
| *Digitipes* | *jangii* | Kali Tiger Reserve, Karwar district, Karnataka, India | 14.9897 | 74.3777 | Joshi & Karanth 2012 |
| *Digitipes* | *jangii* | Kali Tiger Reserve, Karwar district, Karnataka, India | 14.9890 | 74.3712 | Joshi & Karanth 2012 |
| *Digitipes* | *jangii* | Kumta, Uttara Kannada district, Karnataka, India | 14.5163 | 74.5406 | Joshi & Karanth 2012 |
| *Digitipes* | *jangii* | Talacauvery Reserve Forest, Kodagu district, Karnataka, India | 12.3889 | 75.4900 | Joshi & Karanth 2012 |
| *Digitipes* | *jangii* | Kurinjal, Kudremukh National Park, Chikkamagaluru district, India | 13.2018 | 75.1919 | Joshi & Karanth 2012 |
| *Digitipes* | *jangii* | Kurinjal, Kudremukh National Park, Chikkamagaluru district, India | 13.2005 | 75.1929 | Joshi & Edgecombe 2018 |
| *Digitipes* | *jangii* | Tadoli, Kudremukh National Park, Chikkamagaluru district, India | 13.1339 | 75.2782 | Joshi & Karanth 2012 |
| *Digitipes* | *jangii* | Bombay point, Mahabaleshwar Reserve Forest, Satara district, Maharashtra, India | 17.9119 | 73.6368 | Joshi & Karanth 2012 |
| *Digitipes* | *jangii* | Bisale Ghat Reserve Forest, Hassan district, Karnataka, India | 12.7238 | 75.7097 | Joshi & Karanth 2012 |
| *Digitipes* | *jonesii* | Neyyar Wildlife Sanctuary, Thiruvananthapuram district, Kerala, India | 8.6781 | 77.1599 | Joshi & Karanth 2011 |
| *Digitipes* | *jonesii* | Neyyar Wildlife Sanctuary, Thiruvananthapuram district, Kerala, India | 8.6792 | 77.1606 | Joshi & Karanth 2011 |
| *Digitipes* | *jonesii* | Neyyar Wildlife Sanctuary, Thiruvananthapuram district, Kerala, India | 8.6799 | 77.1589 | Joshi & Karanth 2011 |
| *Digitipes* | *jonesii* | Neyyar Wildlife Sanctuary, Thiruvananthapuram district, Kerala, India | 8.6816 | 77.1629 | Joshi & Karanth 2011 |
| *Digitipes* | *jonesii* | Peppara Wildlife Sanctuary, Thiruvananthapuram district, Kerala, India | 8.6663 | 77.1714 | Joshi & Karanth 2011 |
| *Digitipes* | *jonesii* | Peppara Wildlife Sanctuary, Thiruvananthapuram District, Kerala, India | 8.6644 | 77.1788 | Joshi & Karanth, 2011 |
| *Digitipes* | *jonesii* | Peppara Wildlife Sanctuary, Thiruvananthapuram district, Kerala, India | 8.6585 | 77.1779 | Joshi & Karanth 2011 |
| *Digitipes* | *jonesii* | Peppara Wildlife Sanctuary, Thiruvananthapuram District, Kerala, India | 8.6626 | 77.1712 | Joshi & Karanth, 2011 |
| *Digitipes* | *jonesii* | Peppara Wildlife Sanctuary, Thiruvananthapuram district, Kerala, India | 8.6644 | 77.1690 | Joshi & Karanth 2011 |
| *Digitipes* | *jonesii* | Ponmudi Reserve Forest, Thiruvananthapuram district, Kerala, India | 8.7555 | 77.1137 | Joshi & Karanth 2012 |
| *Digitipes* | *jonesii* | Ponmudi Reserve Forest, Thiruvananthapuram district, Kerala, India | 8.7555 | 77.1137 | Joshi, Karanth & Edgecombe, 2020 |
| *Digitipes* | *jonesii* | Ponmudi Reserve Forest, Thiruvananthapuram district, Kerala, India | 8.7562 | 77.1154 | Joshi & Karanth 2012 |
| *Digitipes* | *jonesii* | Pandimatte, Shendurney Wildlife Sanctuary, Kollam district, Kerala, India | 8.8750 | 77.1597 | Joshi & Karanth 2012 |
| *Digitipes* | *jonesii* | Pandimatte, Shendurney Wildlife Sanctuary, Kollam district, Kerala, India | 8.8731 | 77.1628 | Joshi & Karanth 2012 |
| *Digitipes* | *jonesii* | Pandimatte, Shendurney Wildlife Sanctuary, Kollam district, Kerala, India | 8.8732 | 77.1666 | Joshi & Karanth 2012 |
| *Digitipes* | *jonesii* | Pandimatte, Shendurney Wildlife Sanctuary, Kollam district, Kerala, India | 8.8728 | 77.1634 | Joshi & Karanth 2012 |
| *Digitipes* | *jonesii* | Pandimatte, Shendurney Wildlife Sanctuary, Kollam district, Kerala, India | 8.8642 | 77.1797 | Joshi & Karanth 2012 |
| *Digitipes* | *jonesii* | Achankovil Reserve Forest, Pathanamthitta district, Kerala, India | 9.1072 | 77.1306 | Joshi & Karanth 2012 |
| *Digitipes* | *jonesii* | Kottavasal Reserve Forest, Malappuram district, Kerala, India | 9.0703 | 77.2063 | Joshi & Karanth 2012 |
| *Digitipes* | *jonesii* | Kottavasal Reserve Forest, Malappuram district, Kerala, India | 9.0709 | 77.2066 | Joshi & Karanth 2012 |
| *Digitipes* | *jonesii* | Kottavasal Reserve Forest, Malappuram district, Kerala, India | 9.0719 | 77.2060 | Joshi & Karanth 2012 |
| *Digitipes* | *jonesii* | Peppara Wildlife Sanctuary, Thiruvananthapuram district, Kerala, India | 8.6644 | 77.1788 | Joshi & Karanth 2012 |
| *Digitipes* | *jonesii* | Ponmudi, Thiruvananthapuram district, Kerala, India | 8.7300 | 77.1237 | Joshi & Karanth 2012 |
| *Digitipes* | *jonesii* | Eravikulam National Park, Idukki district, Kerala, India | 10.1756 | 77.1062 | Joshi & Karanth 2012 |
| *Digitipes* | *jonesii* | Eravikulam National Park, Idukki district, Kerala, India | 10.2149 | 77.0944 | Joshi & Karanth 2012 |
| *Digitipes* | *jonesii* | Thattekad Bird Sanctuary, Ernakulam district, Kerala, India | 10.1322 | 76.6831 | Joshi & Karanth 2012 |
| *Digitipes* | *jonesii* | Thattekad Bird Sanctuary, Ernakulam district, Kerala, India | 10.1338 | 76.6819 | Joshi & Karanth, 2012 |
| *Digitipes* | *jonesii* | Peechi-Vazhani Wildlife Sanctuary, Thrissur district, Kerala, India | 10.5754 | 76.3854 | Joshi & Karanth 2012 |
| *Digitipes* | *jonesii* | Peechi-Vazhani Wildlife Sanctuary, Thrissur district, Kerala, India | 10.5273 | 76.3500 | Joshi & Karanth 2011 |
| *Digitipes* | *jonesii* | Parambikulam Tiger Reserve, Palakkad district, Kerala, India | 10.3945 | 76.6692 | Joshi & Karanth 2012 |
| *Digitipes* | *jonesii* | Parambikulam Tiger Reserve, Palakkad district, Kerala, India | 10.3798 | 76.6562 | Joshi & Karanth 2012 |
| *Digitipes* | *jonesii* | Sholayar Reserve Forest, Ernakulam district, Kerala, India | 10.3100 | 76.7225 | Joshi & Karanth 2012 |
| *Digitipes* | *jonesii* | Vazhachal Reserve Forest, Thrissur district, Kerala, India | 10.3015 | 76.6053 | Joshi & Karanth 2012 |
| *Digitipes* | *jonesii* | Vellithode, Periyar Tiger Reserve, Kerala, India | 9.4429 | 76.9835 | Joshi & Karanth 2012 |
| *Digitipes* | *jonesii* | Vellithode, Periyar Tiger Reserve, Kerala, India | 9.4406 | 76.9815 | Joshi & Edgecombe 2018 |
| *Digitipes* | *jonesii* | Vellithode, Periyar Tiger Reserve, Kerala, India | 9.4406 | 76.9815 | Joshi & Karanth 2012 |
| *Digitipes* | *jonesii* | Vellithode, Periyar Tiger Reserve, Kerala, India | 9.4430 | 76.9792 | Joshi & Karanth 2012 |
| *Digitipes* | *jonesii* | Vellithode, Periyar Tiger Reserve, Kerala, India | 9.4424 | 76.9831 | Joshi & Karanth 2012 |
| *Digitipes* | *jonesii* | Periyar Tiger Reserve, Pathanamthitta district, Kerala, India | 9.4228 | 76.9894 | Joshi & Karanth 2012 |
| *Digitipes* | *jonesii* | IISER campus, Thiruvananthapuram district, Kerala, India | 8.6827 | 77.1380 | Joshi, Karanth & Edgecombe, 2020 |
| *Digitipes* | *jonesii* | Thenmala, Kollam district, Kerala, India | 8.9417 | 77.1660 | Joshi, Karanth & Edgecombe, 2020 |
| *Digitipes* | *jonesii* | Waynad, Waynad district, Kerala, India | 11.5153 | 76.0395 | Joshi, Karanth & Edgecombe, 2020 |
| *Digitipes* | *nudus* | Silent Valley National Park, Palakkad district, Kerala, India | 11.2089 | 76.4411 | Joshi & Karanth 2011 |
| *Digitipes* | *nudus* | Silent Valley National Park, Palakkad district, Kerala, India | 11.2022 | 76.4392 | Joshi & Karanth 2011 |
| *Digitipes* | *nudus* | Kurinjal, Kudremukh National Park, Chikkamagaluru district, India | 13.2005 | 75.1929 | Joshi & Edgecombe 2018 |
| *Digitipes* | *nudus* | Pambadum Shola National Park, Idukki district, Kerala, India | 10.1756 | 77.1062 | Joshi & Edgecombe 2018 |
| *Digitipes* | *nudus* | Vellimalai, Periyar Tiger Reserve, Idukki district, Kerala, India | 9.5840 | 77.3490 | Joshi & Karanth 2012 |
| *Digitipes* | *nudus* | Vellimalai, Periyar Tiger Reserve, Idukki district, Kerala, India | 9.5286 | 77.3831 | Joshi & Karanth 2012 |
| *Ethmostigmus* | *agasthyamalaiensis* | Parambikulam Tiger Reserve, Palakkad district, Kerala, India | 10.3945 | 76.6692 | Joshi & Edgecombe 2018 |
| *Ethmostigmus* | *agasthyamalaiensis* | Pandimatte, Shendurney Wildlife Sanctuary, Kollam district, Kerala, India | 8.8619 | 77.1795 | Joshi & Edgecombe 2018 |
| *Ethmostigmus* | *agasthyamalaiensis* | Kottavasal Reserve Forest, Malappuram district, Kerala, India | 9.0703 | 77.2063 | Joshi & Edgecombe 2018 |
| *Ethmostigmus* | *agasthyamalaiensis* | Achankovil Reserve Forest, Pathanamthitta district, Kerala, India | 9.1072 | 77.1306 | Joshi & Edgecombe 2018 |
| *Ethmostigmus* | *agasthyamalaiensis* | Pandimatte, Shendurney Wildlife Sanctuary, Kollam district, Kerala, India | 8.8731 | 77.1628 | Joshi & Edgecombe 2018 |
| *Ethmostigmus* | *agasthyamalaiensis* | Neyyar Wildlife Sanctuary, Thiruvananthapuram district, Kerala, India | 8.6781 | 77.1599 | Joshi & Edgecombe 2018 |
| *Ethmostigmus* | *sahyadrensis* | Shirgaokar Road, Amboli, Sindhudurg district, Maharashtra, India | 15.9594 | 73.9997 | Joshi & Karanth 2011 |
| *Ethmostigmus* | *sahyadrensis* | Bombay point, Mahabaleshwar Reserve Forest, Satara district, Maharashtra, India | 17.9119 | 73.6368 | Joshi & Edgecombe 2018 |
| *Ethmostigmus* | *sahyadrensis* | Parikshit Point, Amboli, Sindhudurg district, Maharashtra, India | 15.9450 | 74.0052 | Joshi & Edgecombe 2018 |
| *Ethmostigmus* | *sahyadrensis* | Sadachi Rai, Amboli, Sindhudurg district, Maharashtra, India | 15.9617 | 74.0021 | Joshi & Edgecombe 2018 |
| *Ethmostigmus* | *sahyadrensis* | Lingmala, Mahabaleshwar Reserve Forest, Satara district, Maharashtra, India | 17.9174 | 73.6901 | Joshi & Edgecombe 2018 |
| *Ethmostigmus* | *coonooranus* | Coonoor, Nilgiris district,Tamil Nadu, India | 11.3703 | 76.8041 | Joshi & Edgecombe 2018 |
| *Ethmostigmus* | *coonooranus* | Virajpet, Kodagu district, Karnataka, India | 12.3889 | 75.4900 | Joshi & Karanth 2011 |
| *Ethmostigmus* | *coonooranus* | Parambikulam Tiger Reserve, Palakkad district, Kerala, India | 10.3807 | 76.6576 | Joshi & Edgecombe 2018 |
| *Ethmostigmus* | *coonooranus* | Parambikulam Tiger Reserve, Palakkad district, Kerala, India | 10.4201 | 76.7094 | Joshi & Edgecombe 2018 |
| *Ethmostigmus* | *coonooranus* | Irpu falls, Bramhagiri Wildlife Sanctuary, Kodagu district, Karnataka, India | 12.1700 | 75.6000 | Joshi & Edgecombe 2018 |
| *Ethmostigmus* | *coonooranus* | Coonoor, Nilgiris district, Tamil Nadu, India | 11.3300 | 76.7772 | Jangi & Dass, 1984 |
| *Ethmostigmus* | *praveeni* | Kudremukh National Park, Chikkamagaluru district, Karnataka, India | 13.2018 | 75.1919 | Joshi & Edgecombe 2018 |
| *Ethmostigmus* | *praveeni* | Kudremukh National Park, Chikkamagaluru district, Karnataka, India | 13.1339 | 75.2782 | Joshi & Edgecombe 2018 |
| *Ethmostigmus* | *praveeni* | Sharavathi Wildlife Sanctuary, Shimoga district, Karnataka, India | 14.1158 | 74.7614 | Joshi & Edgecombe 2018 |
| *Ethmostigmus* | *praveeni* | Sakleshpur, Hassan district, Karnataka, India | 12.7238 | 75.7097 | Joshi & Edgecombe 2018 |
| *Ethmostigmus* | *tristis* | Kolli hills, Namakkal district, Tamil Nadu, India | 11.3174 | 78.3471 | Joshi & Edgecombe 2018 |
| *Ethmostigmus* | *tristis* | Kolli hills, Namakkal district, Tamil Nadu, India | 11.3049 | 78.3450 | Joshi & Edgecombe 2018 |
| *Ethmostigmus* | *tristis* | Kolli hills, Namakkal district, Tamil Nadu, India | 11.3583 | 78.3432 | Joshi & Edgecombe 2018 |
| *Ethmostigmus* | *tristis* | Kolli hills, Namakkal district, Tamil Nadu, India | 11.7503 | 78.1885 | Joshi & Edgecombe 2018 |
| *Ethmostigmus* | *tristis* | Kolli hills, Namakkal district, Tamil Nadu, India | 11.3174 | 78.3471 | Joshi & Edgecombe 2018 |
| *Ethmostigmus* | *tristis* | Kolli hills, Namakkal district, Tamil Nadu, India | 11.3247 | 78.3414 | Joshi & Edgecombe 2018 |
| *Ethmostigmus* | *tristis* | Yercaud, Salem district, Tamil Nadu, India | 11.7763 | 78.1862 | Joshi & Edgecombe 2018 |
| *Ethmostigmus* | *tristis* | Yercaud, Salem district, Tamil Nadu, India | 11.7553 | 78.2000 | Joshi & Edgecombe 2018 |
| *Rhysida* | *konda* | Sitakund, Mayurbhanj district, Odisha, India | 21.9200 | 86.5600 | Joshi & Edgecombe 2018 |
| *Rhysida* | *konda* | Bali, Mayurbhanj district, Odisha, India | 21.0004 | 84.6965 | Joshi & Edgecombe 2018 |
| *Rhysida* | *konda* | Devmali, Koraput district, Odisha, India | 18.6600 | 82.9900 | Joshi & Edgecombe 2018 |
| *Rhysida* | *konda* | Gudem, Visakhapatnam district, Andhra Pradesh, India | 18.1862 | 83.1482 | Joshi, Karanth & Edgecombe, 2020 |
| *Rhysida* | *aspinosa* | Neyyar Wildlife Sanctuary, Thiruvananthapuram district, Kerala, India | 8.6799 | 77.1589 | Joshi & Edgecombe 2018 |
| *Rhysida* | *aspinosa* | Rockwood estate, Shendurney Wildlife Sanctuary, Kollam district, Kerala, India | 8.8751 | 77.1186 | Joshi & Edgecombe 2018 |
| *Rhysida* | *aspinosa* | Achankovil Reserve Forest, Pathanamthitta district, Kerala, India | 9.1269 | 77.1806 | Joshi & Edgecombe 2018 |
| *Rhysida* | *aspinosa* | Periyar Tiger Reserve, Idukki district, Kerala, India | 9.5755 | 77.3362 | Joshi & Edgecombe 2018 |
| *Rhysida* | *crassispina* | Matheran, Raigad district, Maharashtra, India | 18.9893 | 73.2570 | Joshi & Edgecombe 2018 |
| *Rhysida* | *crassispina* | Matheran, Raigad district, Maharashtra, India | 18.9616 | 73.2660 | Joshi & Edgecombe 2018 |
| *Rhysida* | *crassispina* | Matheran, Raigad district, Maharashtra, India | 18.9904 | 73.2878 | Joshi, Karanth & Edgecombe, 2020 |
| *Rhysida* | *crassispina* | Matheran, Raigad district, Maharashtra, India | 19.0111 | 73.2845 | Jangi & Dass, 1984 |
| *Rhysida* | *lewisi* | Anshi-Dandeli Tiger Reserve, Karwar district, Karnataka, India | 14.9890 | 74.3712 | Joshi & Karanth 2011 |
| *Rhysida* | *lewisi* | Kumta, Uttara Kannada district, Karnataka, India | 14.5257 | 74.6025 | Joshi & Karanth 2011 |
| *Rhysida* | *lewisi* | Radhanagari Wildlife Sanctuary, Kolhapur district, Maharashtra, India | 16.2917 | 73.9083 | Joshi & Karanth 2011 |
| *Rhysida* | *lewisi* | Kurinjal, Kudremukh National Park, Chikkamagaluru district, India | 13.2018 | 75.1919 | Joshi & Edgecombe 2018 |
| *Rhysida* | *lewisi* | Tadoli, Kudremukh National Park, Chikkamagaluru district, India | 13.1339 | 75.2782 | Joshi & Edgecombe 2018 |
| *Rhysida* | *lewisi* | Amboli, Sindhudurg district, Maharashtra, India | 15.9594 | 73.9997 | Joshi & Edgecombe 2018 |
| *Rhysida* | *lewisi* | Kudremukh National Park, Chikkamagaluru district, Karnataka, India | 13.1341 | 75.3154 | Joshi & Edgecombe 2018 |
| *Rhysida* | *lewisi* | Talacauvery, Kodagu district, Karnataka, India | 12.4055 | 75.5199 | Joshi & Edgecombe 2018 |
| *Rhysida* | *longipes* | Chatancode, Vidura, Thiruvananthapuram district, Kerala, India | 8.6622 | 77.1509 | Joshi & Edgecombe 2018 |
| *Rhysida* | *longipes* | Indian Institute of Science Campus, Bangalore district, Karnataka, India | 13.0238 | 77.5671 | Joshi & Edgecombe 2018 |
| *Rhysida* | *longipes* | Dimbam Ghat Road, Erode district, Tamil Nadu, India | 11.6344 | 77.1286 | Joshi & Edgecombe 2018 |
| *Rhysida* | *longipes* | Matheran, Raigad district, Maharashtra, India | 18.9900 | 73.2400 | Joshi & Edgecombe 2018 |
| *Rhysida* | *longipes* | Mysore district, Karnataka, India | 12.2700 | 76.2700 | Joshi & Edgecombe 2018 |
| *Rhysida* | *longipes* | Bondla, North Goa district, Goa, India | 15.4401 | 74.1053 | Jangi & Dass, 1984 |
| *Rhysida* | *longipes* | Amravati, Amravati district, Maharashtra, India | 20.9342 | 77.7277 | Jangi & Dass, 1984 |
| *Rhysida* | *longipes* | Nagpur, Nagpur district, India | 21.1813 | 79.0188 | Jangi & Dass, 1984 |
| *Rhysida* | *longipes* | Kumta, Uttara Kannada district, India | 14.4355 | 74.4261 | Jangi & Dass, 1984 |
| *Rhysida* | *pazhuthara* | Peppara Wildlife Sanctuary, Thiruvananthapuram district, Kerala, India | 8.6644 | 77.1788 | Joshi & Karanth 2011 |
| *Rhysida* | *pazhuthara* | Ponmudi, Thiruvananthapuram district, Kerala, India | 8.7300 | 77.1237 | Joshi & Karanth 2011 |
| *Rhysida* | *pazhuthara* | Rockwood estate, Shendurney Wildlife Sanctuary, Kollam district, Kerala, India | 8.8751 | 77.1186 | Joshi & Edgecombe 2018 |
| *Rhysida* | *pazhuthara* | Rosemla forest, Shendurney Wildlife Sanctuary, Kollam district, Kerala, India | 8.9535 | 77.1786 | Joshi & Edgecombe 2018 |
| *Rhysida* | *pazhuthara* | Pandimatte, Shendurney Wildlife Sanctuary, Kollam district, Kerala, India | 8.8732 | 77.1666 | Joshi & Edgecombe 2018 |
| *Rhysida* | *sada* | Bhimashankar, Pune district, Maharashtra, India | 19.0777 | 73.5505 | Joshi & Edgecombe 2018 |
| *Rhysida* | *sada* | Phansad, Raigad district, Maharashtra, India | 18.4377 | 72.9355 | Joshi & Edgecombe 2018 |
| *Rhysida* | *sada* | Phansad, Raigad district, Maharashtra, India | 18.4152 | 72.9712 | Joshi & Edgecombe 2018 |
| *Rhysida* | *sada* | Bhimashankar, Pune district, Maharashtra, India | 19.1323 | 73.5701 | Joshi & Edgecombe 2018 |
| *Rhysida* | *sada* | Phansad, Raigad district, Maharashtra, India | 18.4270 | 72.9563 | Joshi, Karanth & Edgecombe, 2020 |
| *Rhysida* | *sada* | Bhimashankar, Pune district, Maharashtra, India | 19.0633 | 73.5396 | Joshi, Karanth & Edgecombe, 2020 |
| *Rhysida* | *trispinosa* | Ramanagar, Ramanagara district, Karnataka, India | 15.4090 | 74.4762 | Joshi & Karanth 2011 |
| *Rhysida* | *trispinosa* | IISc Campus, Bangalore district, Karnataka, India | 13.0238 | 77.5671 | Joshi & Karanth 2011 |
| *Rhysida* | *trispinosa* | Neyyar Wildlife Sanctuary, Thiruvananthapuram district, Kerala, India | 8.6799 | 77.1589 | Joshi & Karanth 2011 |
| *Rhysida* | *trispinosa* | Devarayanadurga, Tumkur district, Karnataka, India | 13.0381 | 77.2001 | Joshi & Karanth 2011 |
| *Rhysida* | *trispinosa* | Castle-rock, Uttara Kannada district, Karnatka, India | 15.3698 | 74.3063 | Joshi & Edgecombe 2018 |
| *Rhysida* | *trispinosa* | Ranebennur, Haveri district, Karnataka, India | 14.9217 | 75.2480 | Joshi & Karanth 2011 |
| *Rhysida* | *trispinosa* | Biligirirangana Hills, Chamarajanagar district, Karnataka, India | 11.9052 | 77.1597 | Joshi & Edgecombe 2018 |
| *Rhysida* | *trispinosa* | Thattekad Bird Sanctuary, Ernakulam district, Kerala, India | 10.1322 | 76.6831 | Joshi & Edgecombe 2018 |
| *Rhysida* | *trispinosa* | Kangundi, Chittoor district, Andhra Pradesh, India | 12.7686 | 78.4337 | Joshi & Edgecombe 2018 |
| *Rhysida* | *trispinosa* | Nayanoor, Chittoor district, Andhra Pradesh, India | 12.6910 | 78.4360 | Joshi & Edgecombe 2018 |
| *Rhysida* | *trispinosa* | Achankovil Reserve Forest, Pathanamthitta district, Kerala, India | 9.1072 | 77.1306 | Joshi & Edgecombe 2018 |
| *Rhysida* | *trispinosa* | Javadi hills, Tiruvannamalai district, Tamil Nadu, India | 12.6005 | 78.8447 | Joshi & Edgecombe 2018 |
| *Rhysida* | *trispinosa* | Chinnar Wildlife Sanctuary, Idukki district, Kerala, India | 10.3067 | 77.2060 | Joshi & Edgecombe 2018 |
| *Rhysida* | *trispinosa* | Srisailam, Kurnool district, Andhra Pradesh, India | 15.9790 | 78.8480 | Joshi & Edgecombe 2018 |
| *Rhysida* | *trispinosa* | Palaruvi Waterfalls, Kollam district, Kerala, India | 8.9417 | 77.1660 | Joshi, Karanth & Edgecombe, 2020 |
| *Rhysida* | *trispinosa* | Kanakapura, Bangalore district, Karnataka, India | 12.8316 | 77.5006 | Joshi, Karanth & Edgecombe, 2020 |
| *Rhysida* | sp. 1 | Talacauvery, Kodagu district, Karnataka, India | 12.4055 | 75.5199 | Joshi & Edgecombe 2018 |
| *Rhysida* | sp. 1 | Siruvani Reserve Forest, Palakkad district, Kerala, India | 10.9686 | 76.6669 | Joshi & Edgecombe 2018 |
| *Rhysida* | sp. 1 | Javadi hills, Tiruvannamalai district, Tamil Nadu, India | 12.6001 | 78.8369 | Joshi & Edgecombe 2018 |
