## Supporting Information Appendix S2 for "Spatial patterns of phylogenetic diversity and endemism in the Western Ghats, India: a case study using ancient predatory arthropods"

**Appendix S2.** Description of Maxent species distribution modeling approach and results.

**Maxent modeling approach and evaluation**

**Sampling bias**

With unequal sampling effort in a complex landscape with strong latitudinal and elevation gradients, randomly picked background locations can lead to a poor estimate of environmental variation in the model extent, biasing predictions towards sampled environments rather than capturing species preferences (Phillips et al., 2009; Elith et al., 2011). To correct for this, we built a model of sampling bias with all surveyed locations and used its predictions (Fig. S2.1) to weigh the selection of background locations for species distribution models (following Phillips et al., 2009). The same set of background locations was used across all species being modelled.

**Predictor variables**

We used all 19 bioclimatic variables and the elevation layer from the WorldClim database (Fick & Hijmans, 2017) at 30 arcsecond resolution (~1 km at the equator). These variables were derived from monthly data on temperature, precipitation, solar radiation, wind speed and water vapour pressure, summarized over the years 1971-2000, and elevation was derived from SRTM data (http://srtm.csi.cgiar.org/). We additionally used soil type (ATREE Spatial Archive, 2020) as a predictor variable which might influence the distribution of centipedes which are soil arthropods. All of these environmental variables were used as predictors, allowing the regularization procedure in the Maxent algorithm to select features that are important to habitat suitability (Elith et al., 2011).

**Maxent parameters**

We used default settings for feature classes (linear, quadratic, product, hinge, threshold), which are selected by the algorithm based on the number of presence locations available in each model, and default regularization parameters. Model predictions were obtained as maps of the logistic output (ranging from 0 to 1) provided by Maxent (Phillips & Dudik, 2008), which can be interpreted as a measure of relative habitat suitability for a species in each cell of the model extent (Phillips et al., 2006; Elith et al., 2011).

**Model evaluation**

We calculated Area Under the Receiver Operator Curve (AUC; Fielding & Bell, 1998) and the True Skill Statistic (TSS; Allouche et al., 2006) for each model, which assess model accuracy based on sensitivity (true positive rate) and specificity (true negative rate). AUC is derived using a range of thresholds over model predictions to obtain a binary presence-absence map in each instance, which is used to calculate specificity and sensitivity. It ranges from 0 to 1, with a value of 0.5 indicating a model that is no better than random. It can be interpreted as the probability that a model assigns a greater value of habitat suitability to a randomly picked presence location as compared to a randomly picked background location (Merow et al., 2013). TSS is based on a single threshold to convert model predictions into a binary map (Allouche et al., 2006) and we used a threshold that gives the maximum sum of sensitivity and specificity (Liu et al, 2005). TSS ranges from -1 to +1, with a value of 0 indicating that a model that is no better than random.

We estimated error in model AUC and TSS, by five-fold cross-validation of species data-sets which had more than five presence locations. Presence locations were split into five subsets, where four subsets were used for model training and the fifth was withheld for model evaluation. Since most species had few data points, we also performed a jackknifing test for small sample sizes (Pearson et al., 2007). Models were built leaving out each presence location in turn, and evaluated based on accurate assignments of the dropped presence location. These results are used to assess if model predictions are significantly different from random assignments.

**Caveats**

**Sample size**

In an assessment of performance of species distribution models (SDMs) across sample sizes, Wisz et al. (2008) used a range of input data, the lowest being 30 presence locations, and found that Maxent performs more consistently across sample sizes as compared to other modeling algorithms. Though all of the centipede species modeled in our study have fewer point locations than this, with a minimum of three presence locations, other tests with small sample sizes have shown that Maxent models can provide useful predictions with even 5-10 presence locations. These tests also indicated that prediction accuracy improved when the small range sizes of species are related to strong environmental constraints (Hernandez et al., 2006).

Despite using different methods of model evaluation (AUC, TSS and Jack-knifing test for small sample sizes), which detect if the model is better than random, it is hard to estimate the range of error (underfitting or overfitting) in our models given the lack of absence data in our study. Given the nature of species biology, where centipedes often have narrow distributions and are sparsely distributed within their range, this is an unavoidable problem. However, we believe that these species-specific model predictions would be helpful in surveying potential areas of distribution, which can add to the density of presence locations in the future.

**Number of environmental predictors**

We used a wide suite of available predictor variables for our models as there might be interspecific differences in tolerance to environmental variables, especially given the wide variation in extent of distribution between species of interest. Therefore, omitting variables that may be suitable for a subset of species can lead to an increase in prediction error (Arau ́jo & Peterson, 2012). Though decrease in prediction accuracy with the number of predictor variables has been found to be less of a problem with complex SDM algorithms such as Maxent (Júnior & Nóbrega, 2018), it is hard to perform rigorous tests of accuracy without presence-absence test data (Phillips et al., 2009).

The regularization procedure within the Maxent algorithm prevents model overfitting, which reduces the need to select a subset of uncorrelated predictor variables for the modeling exercise, though some level of selection based on relevance is recommended (Elith et al., 2011). However, other recommendations suggest that screening of relevant predictor variables must be done in a similar manner to regression based analyses (Merow et al., 2013).

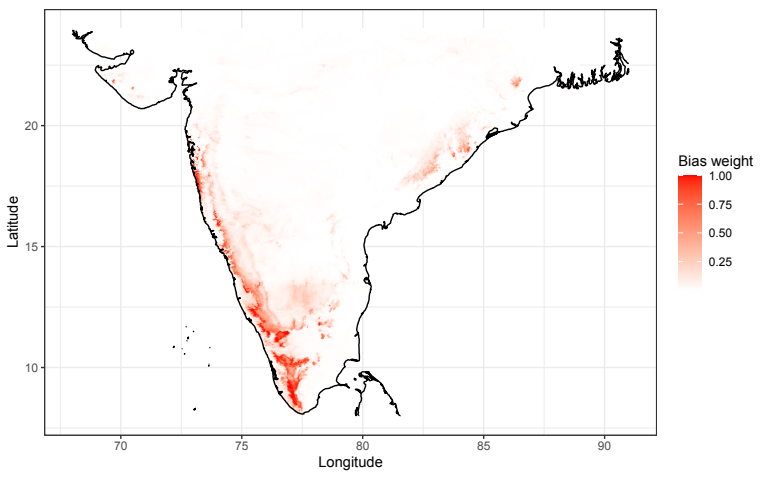
**Figure S2.1.** Prediction map of the sampling bias model which was used to weight selection of background locations for the Maxent species distribution models.

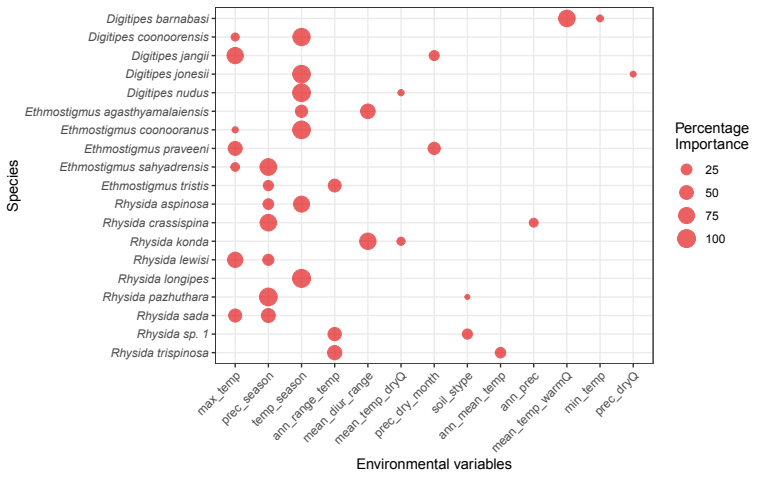
**Figure S2.2.** Permutation importance of variables used to run Maxent species distribution models for each species. Variables with the highest two values of permutation importance are indicated using the position of the filled circle on the x-axis, while the size of the circle corresponds to the magnitude of percentage permutation importance.

**Variable codes:**

**max_temp** – maximum temperature of the warmest month, **prec_season** – precipitation seasonality, **temp_season** – temperature seasonality, **ann_range_temp** – annual range of temperature, **mean_diur_range** – mean diurnal range of temperature, **mean_temp_dryQ** – mean temperature of the driest quarter, **prec_dry_month** – precipitation of the driest month, **soil_stype** – soil sub-type, **ann_mean_temp** – annual mean temperature, **ann_prec** – annual precipitation, **mean_temp_warmQ** – mean temperature of the warmest quarter, **min_temp** – minimum temperature of the coldest month, **prec_dryQ** – preciptation of the driest quarter.

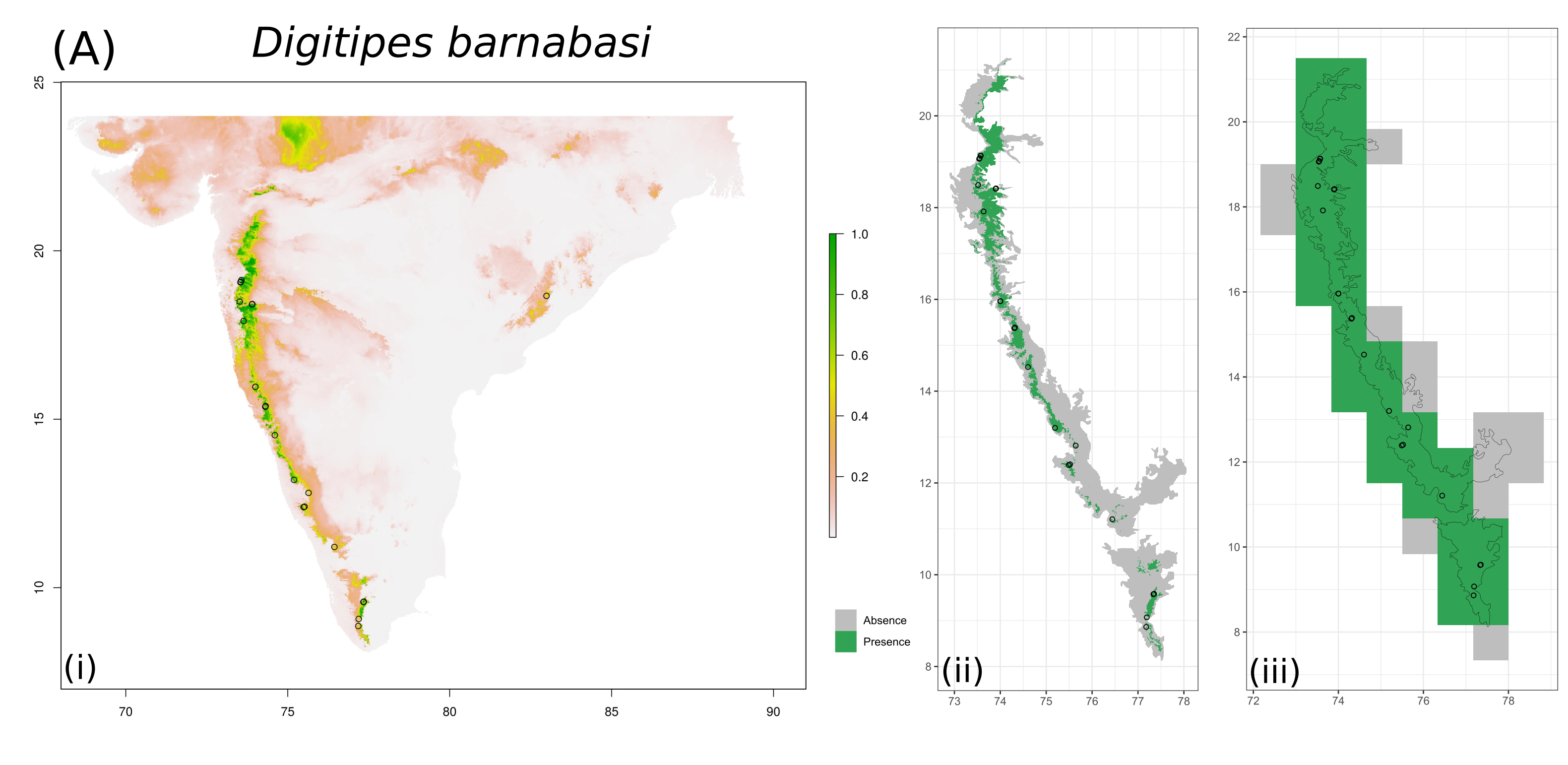

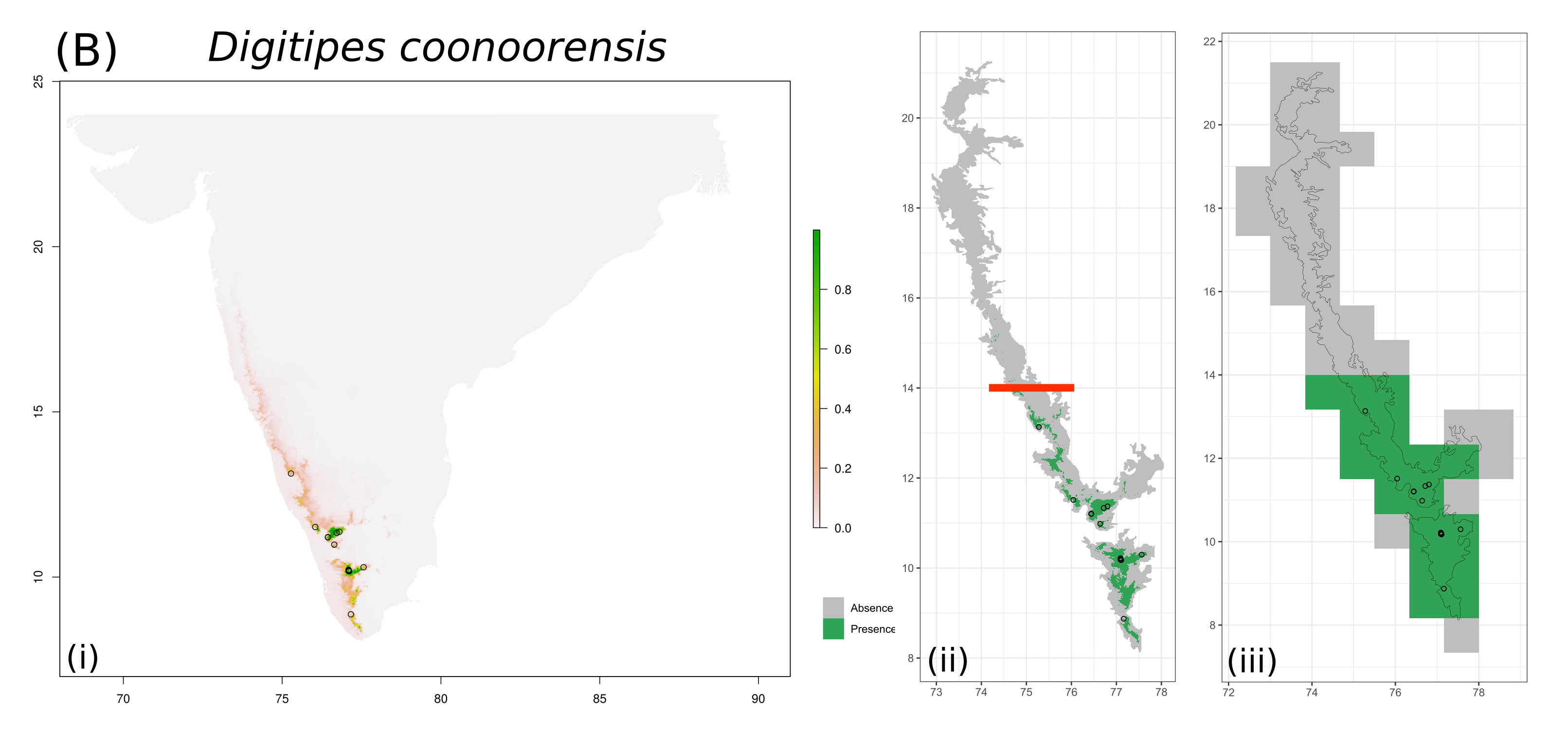

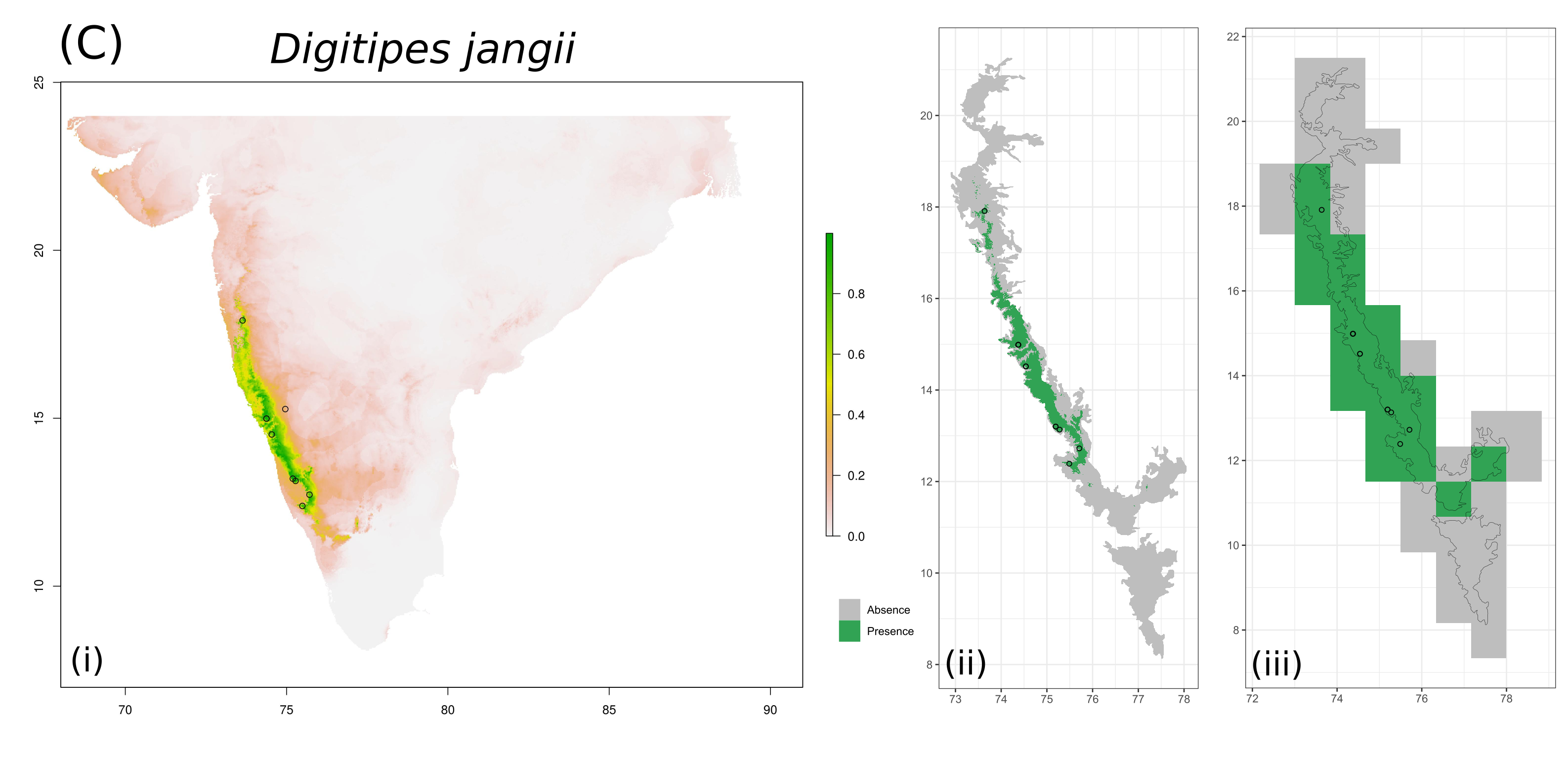

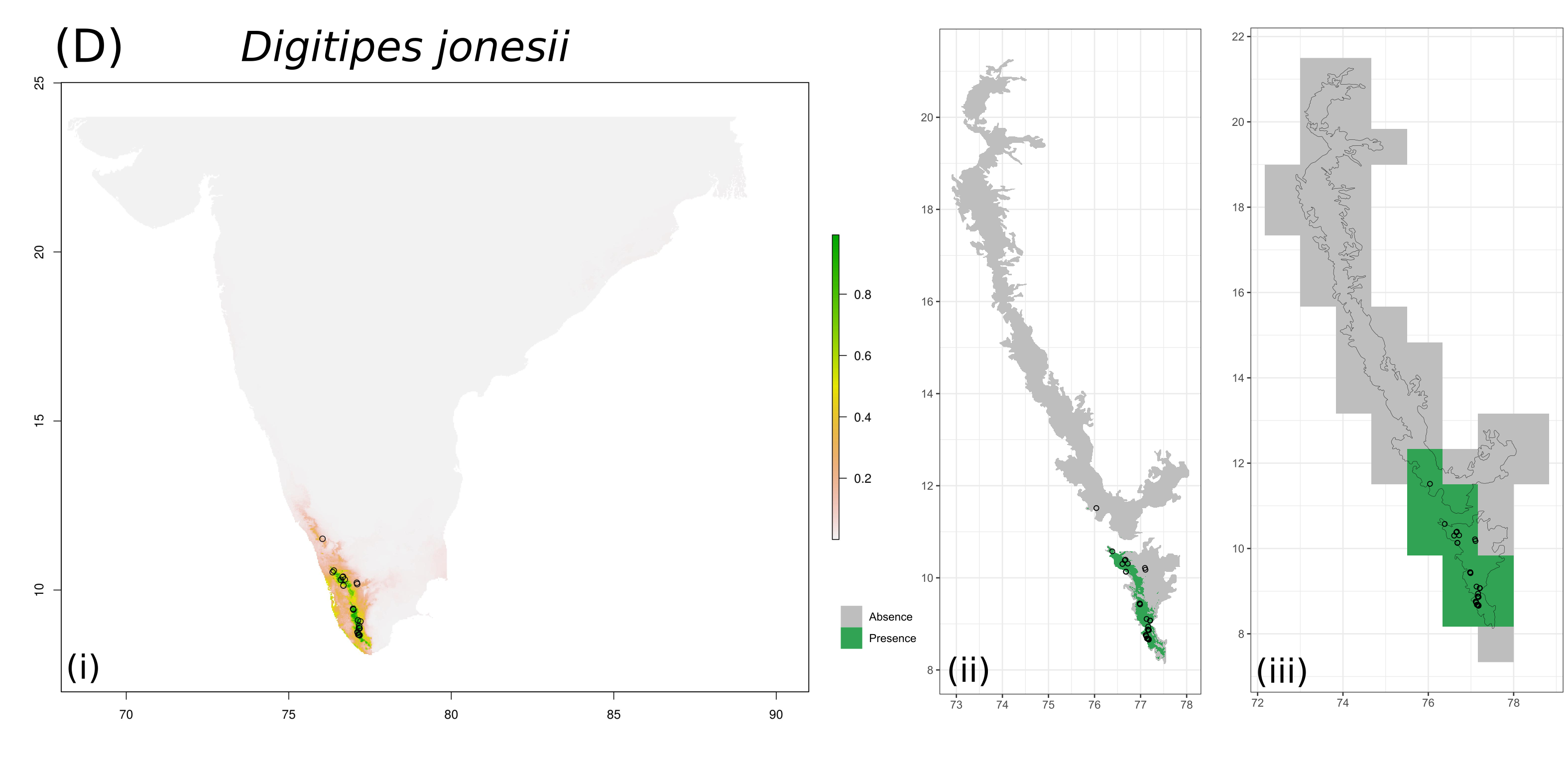

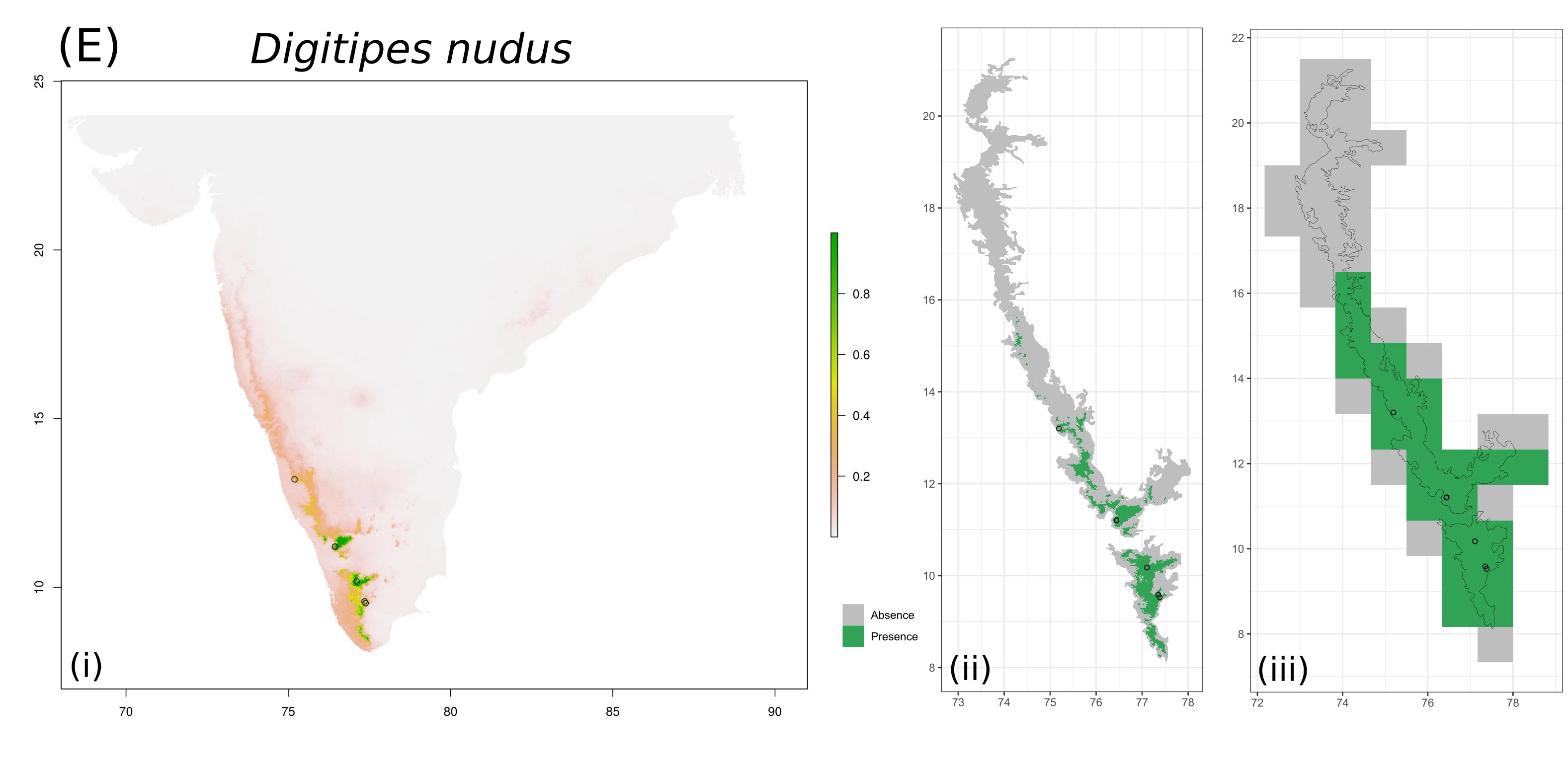

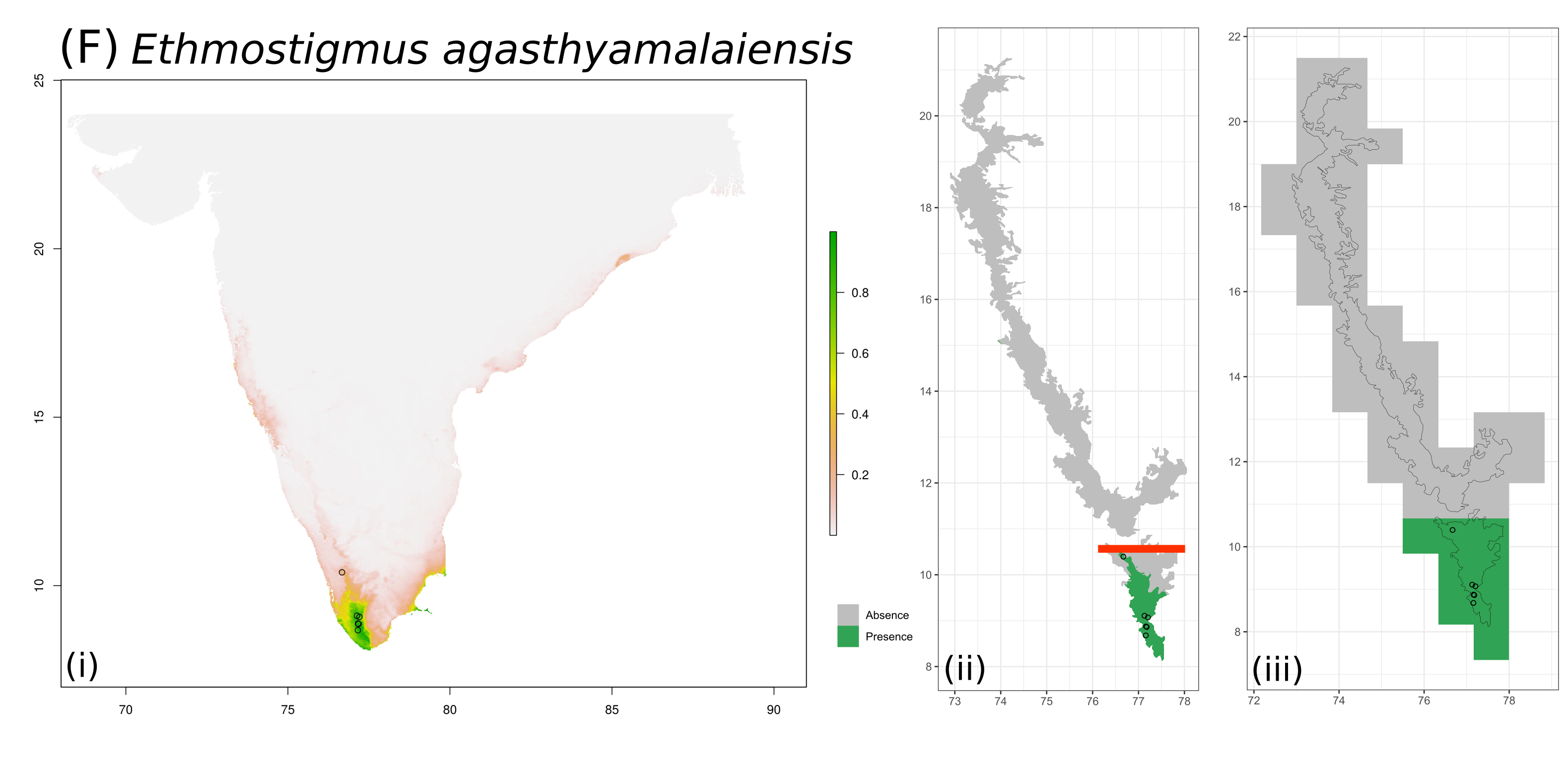

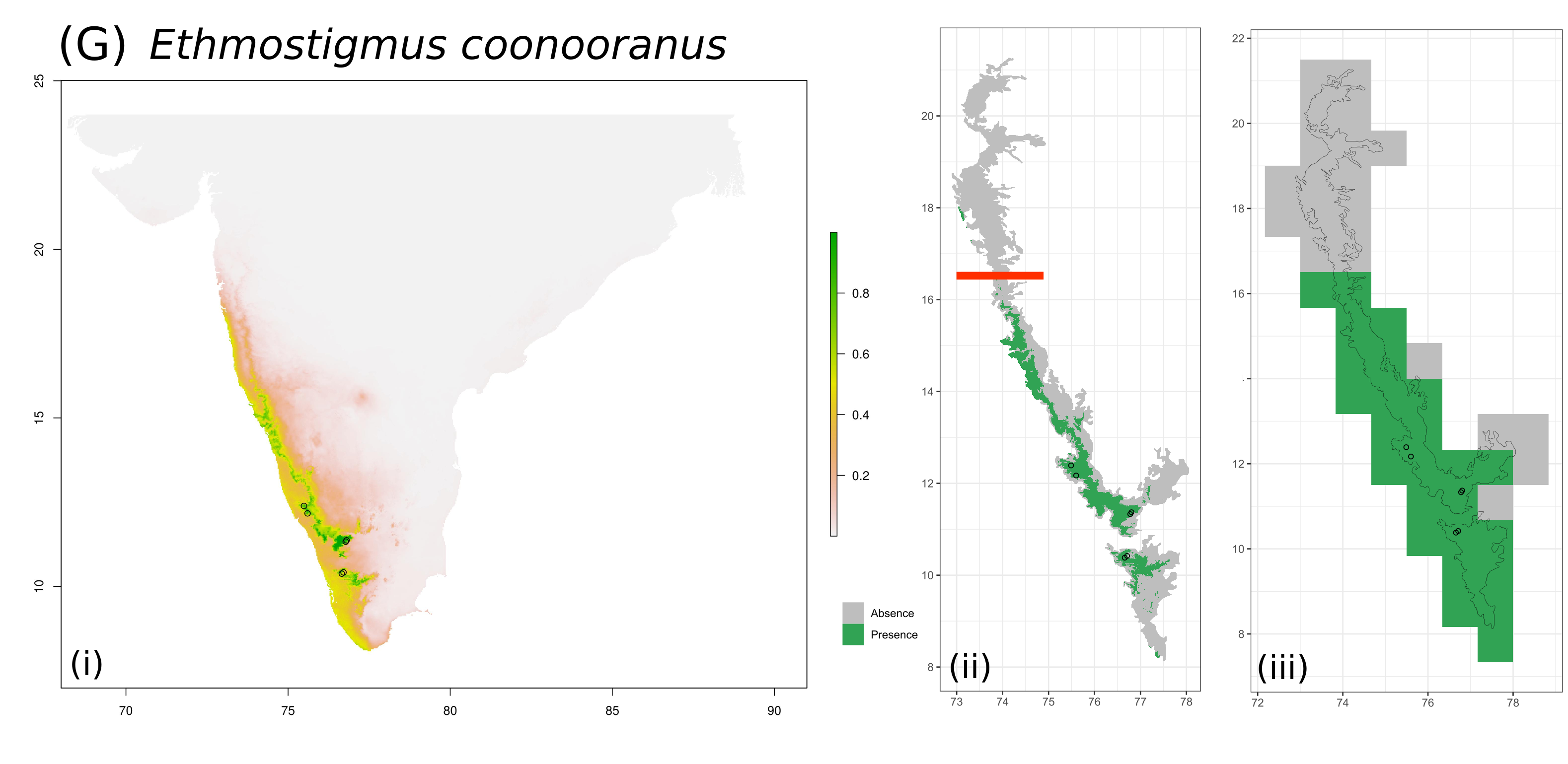

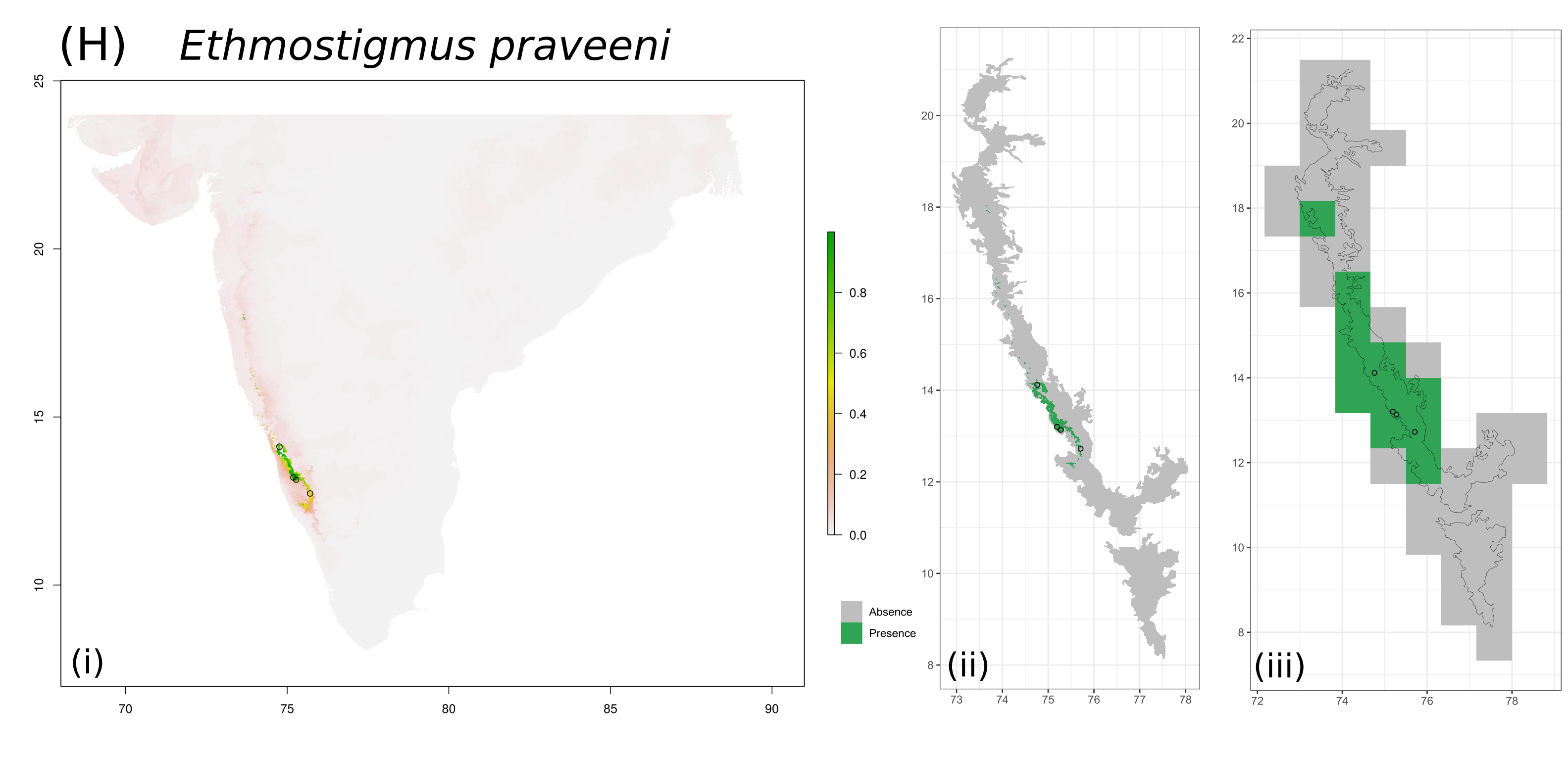

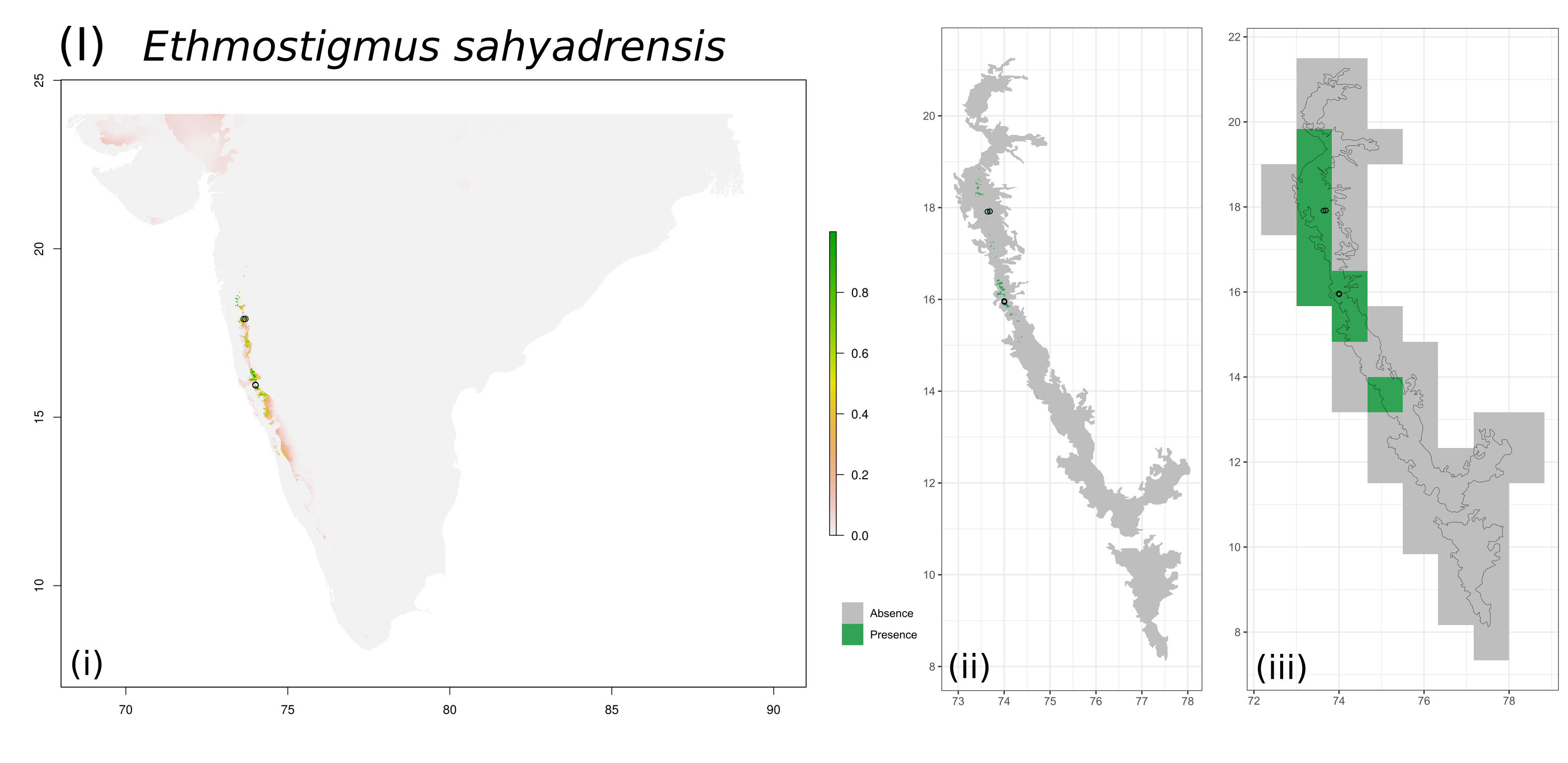

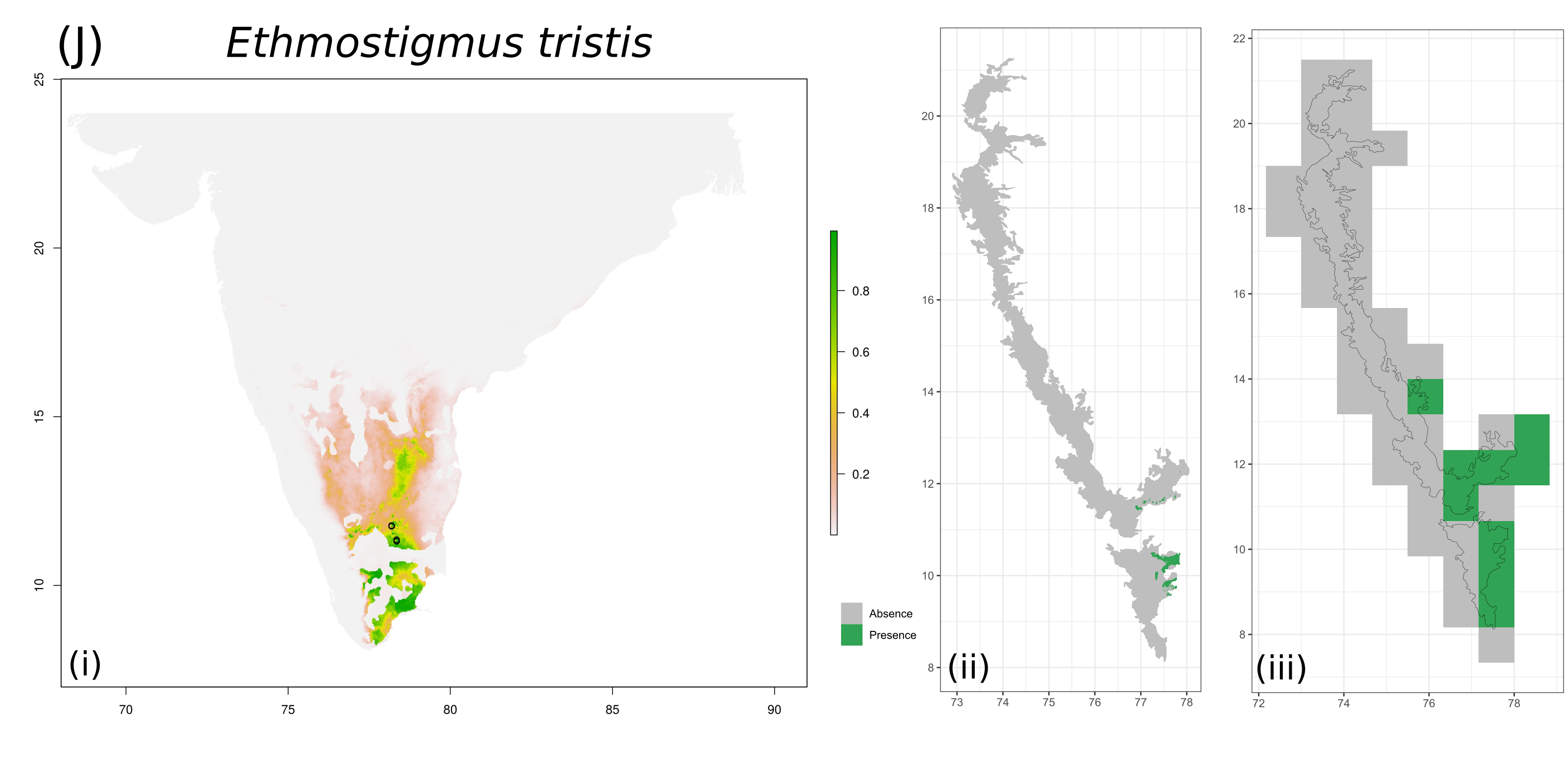

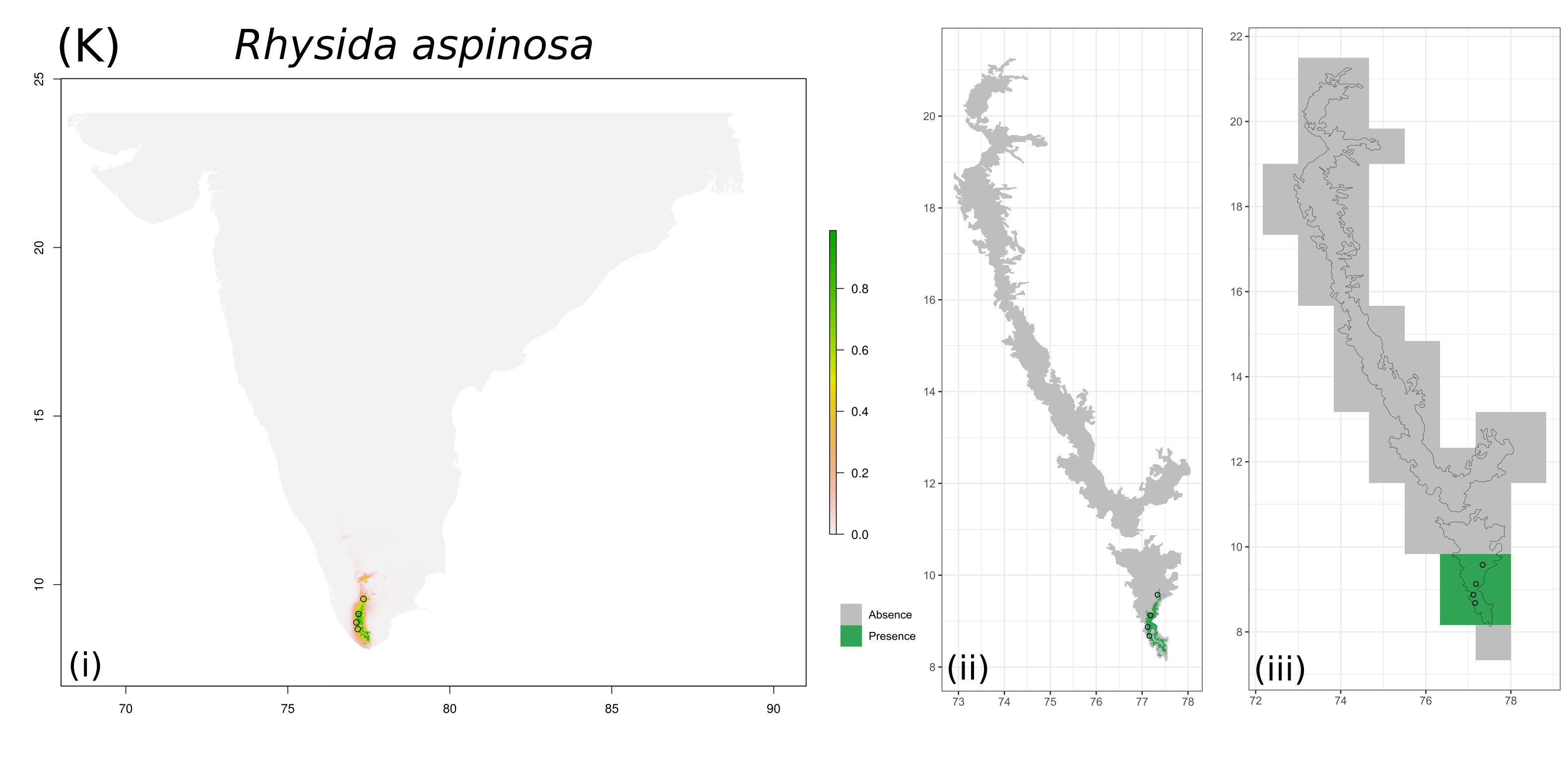

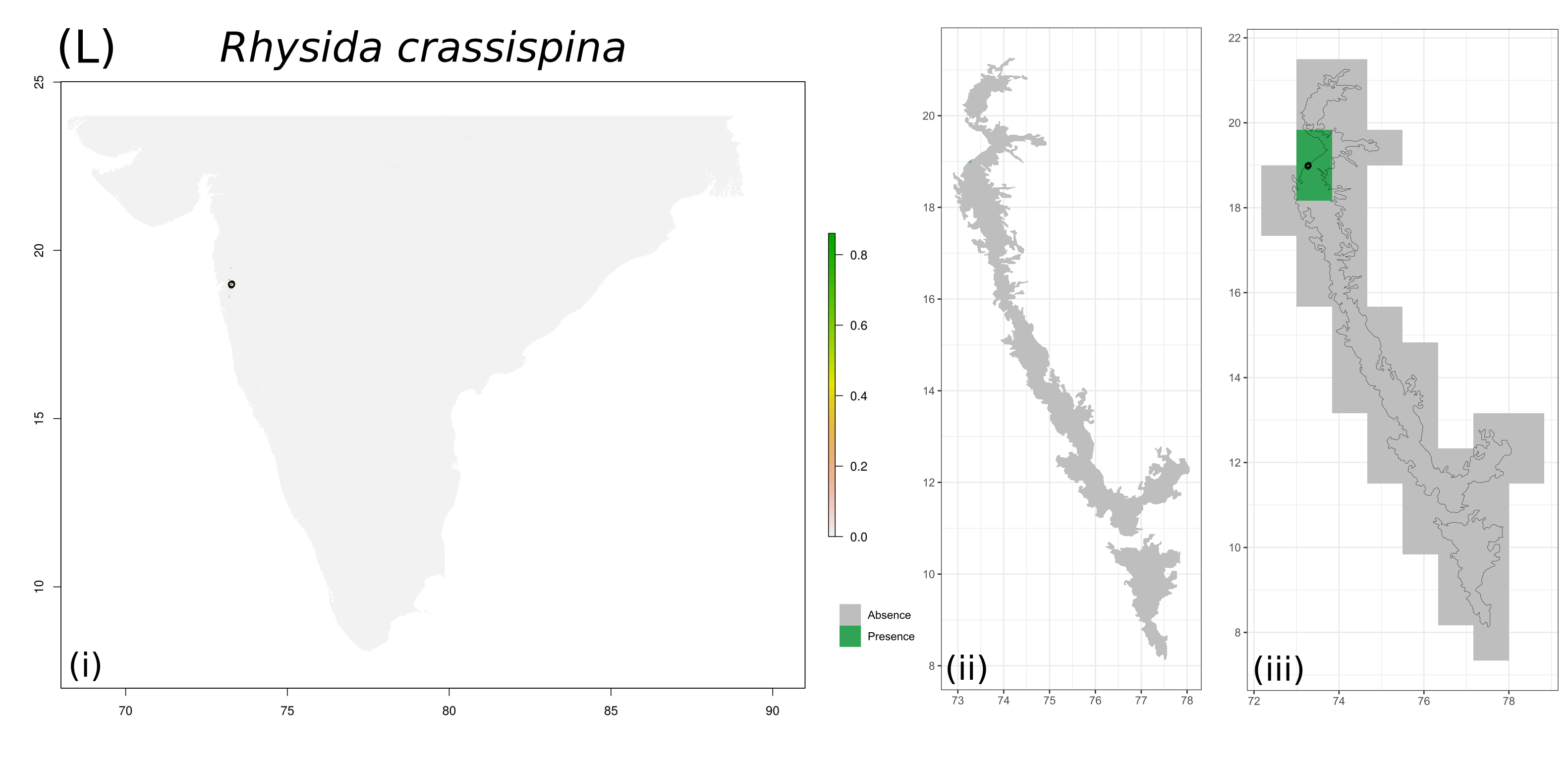

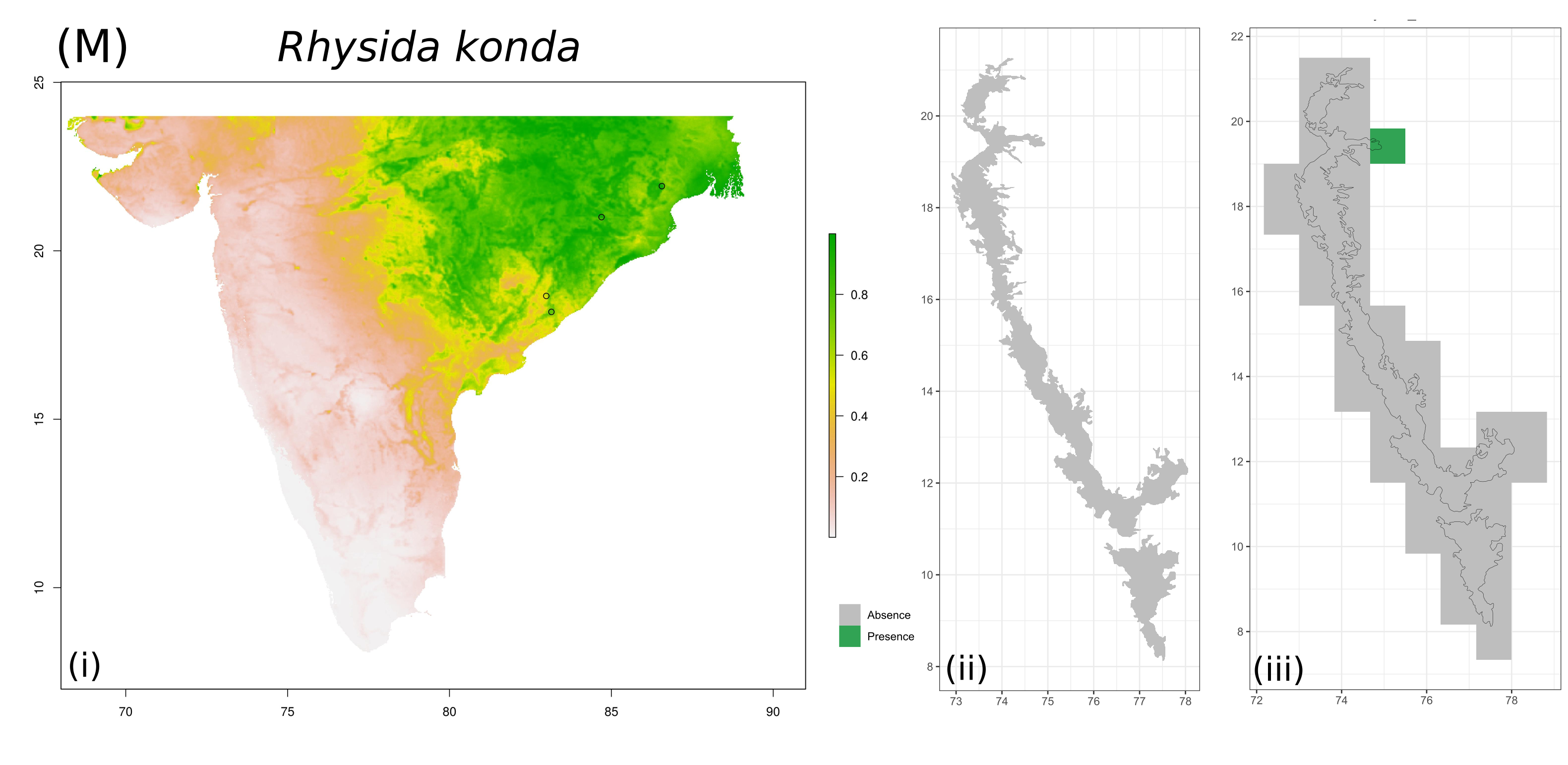

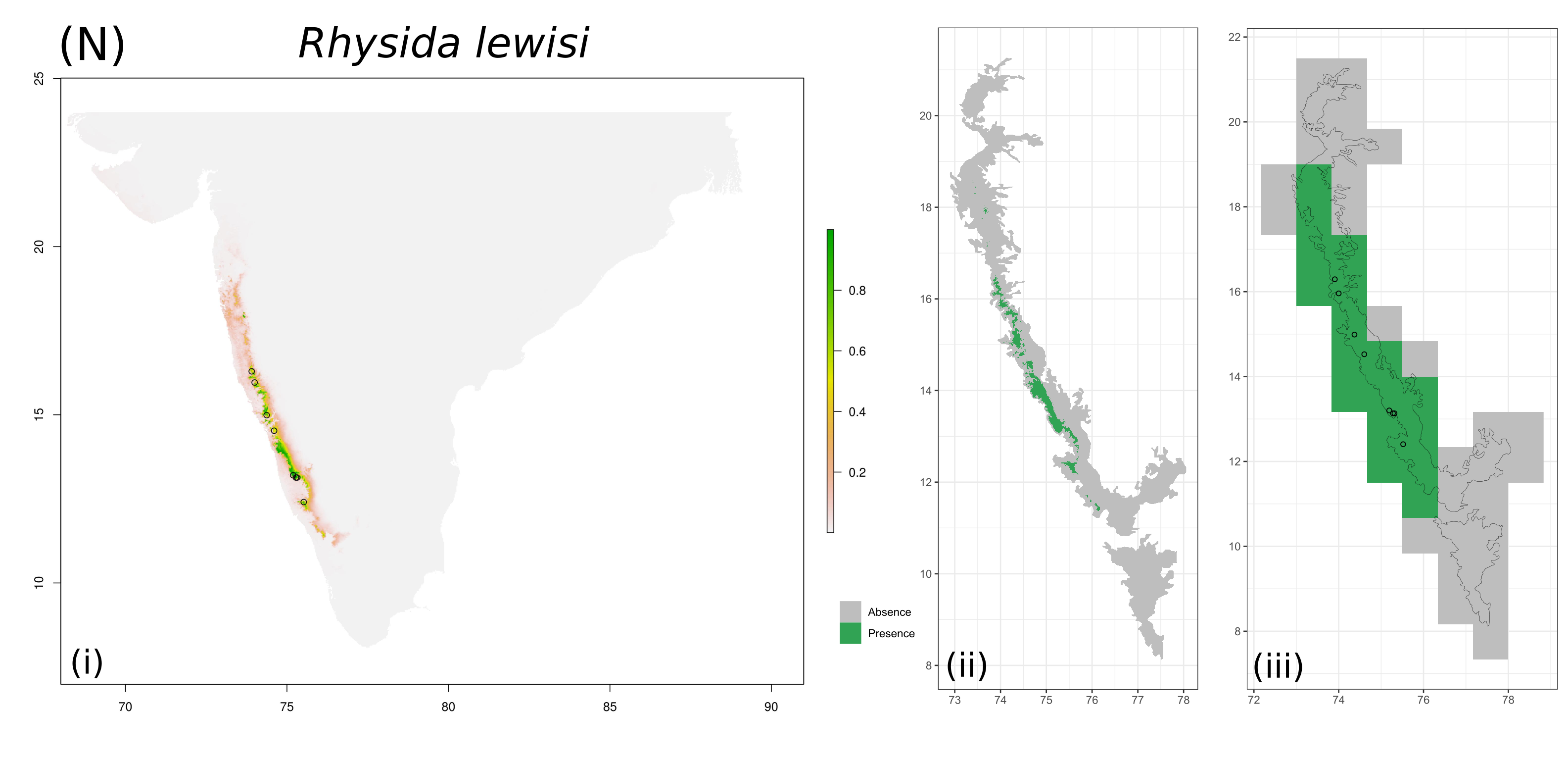

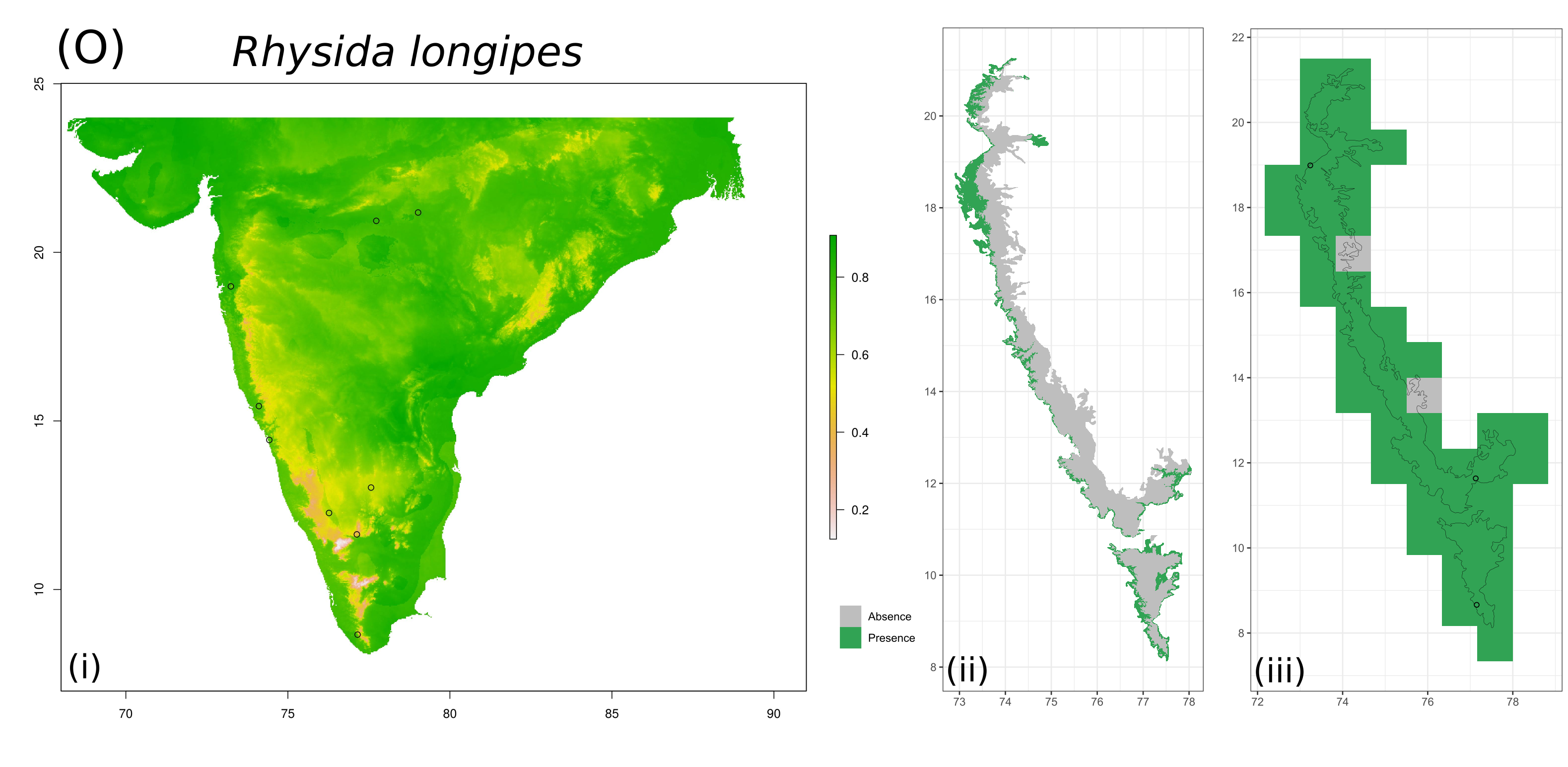

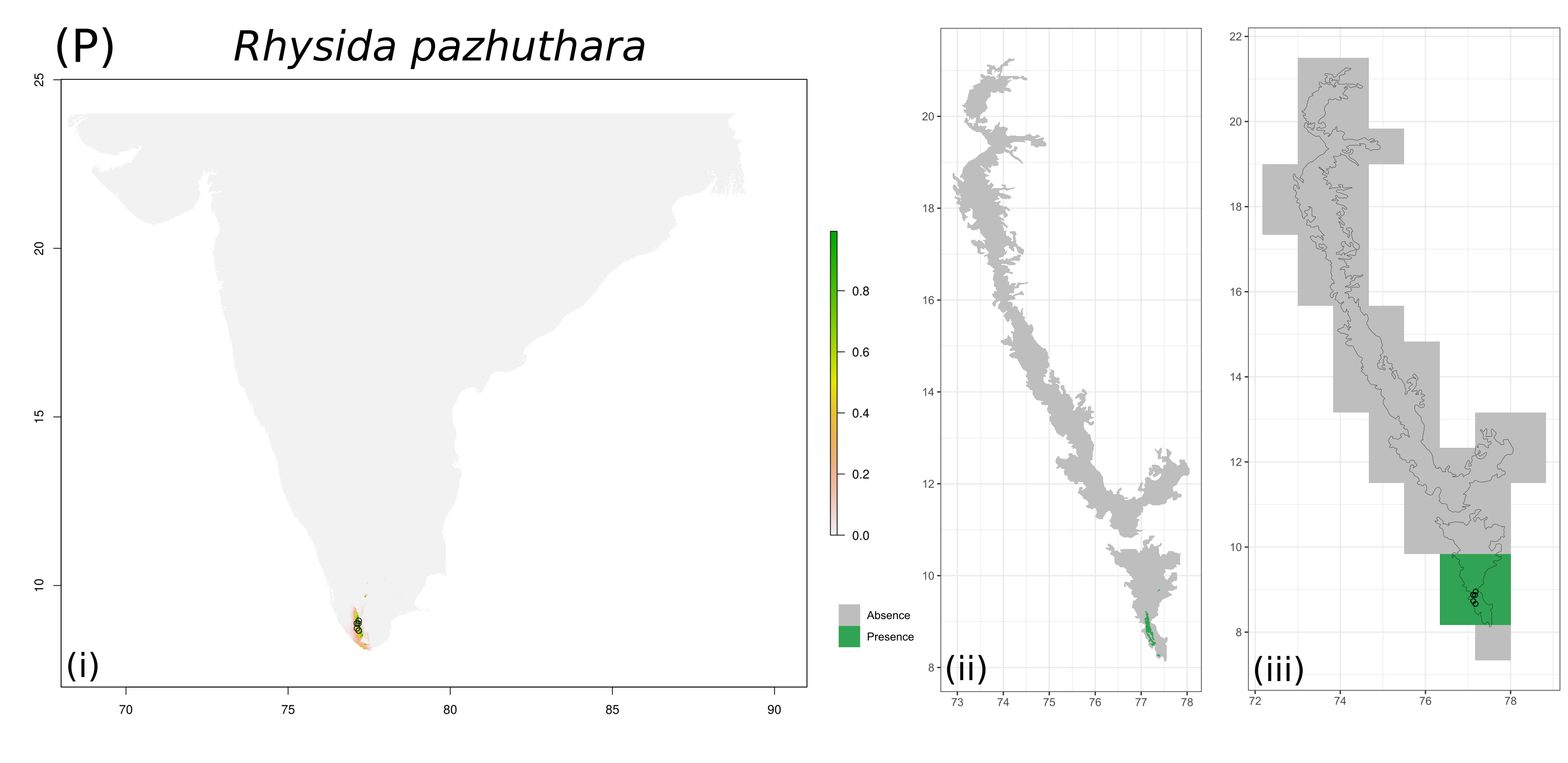

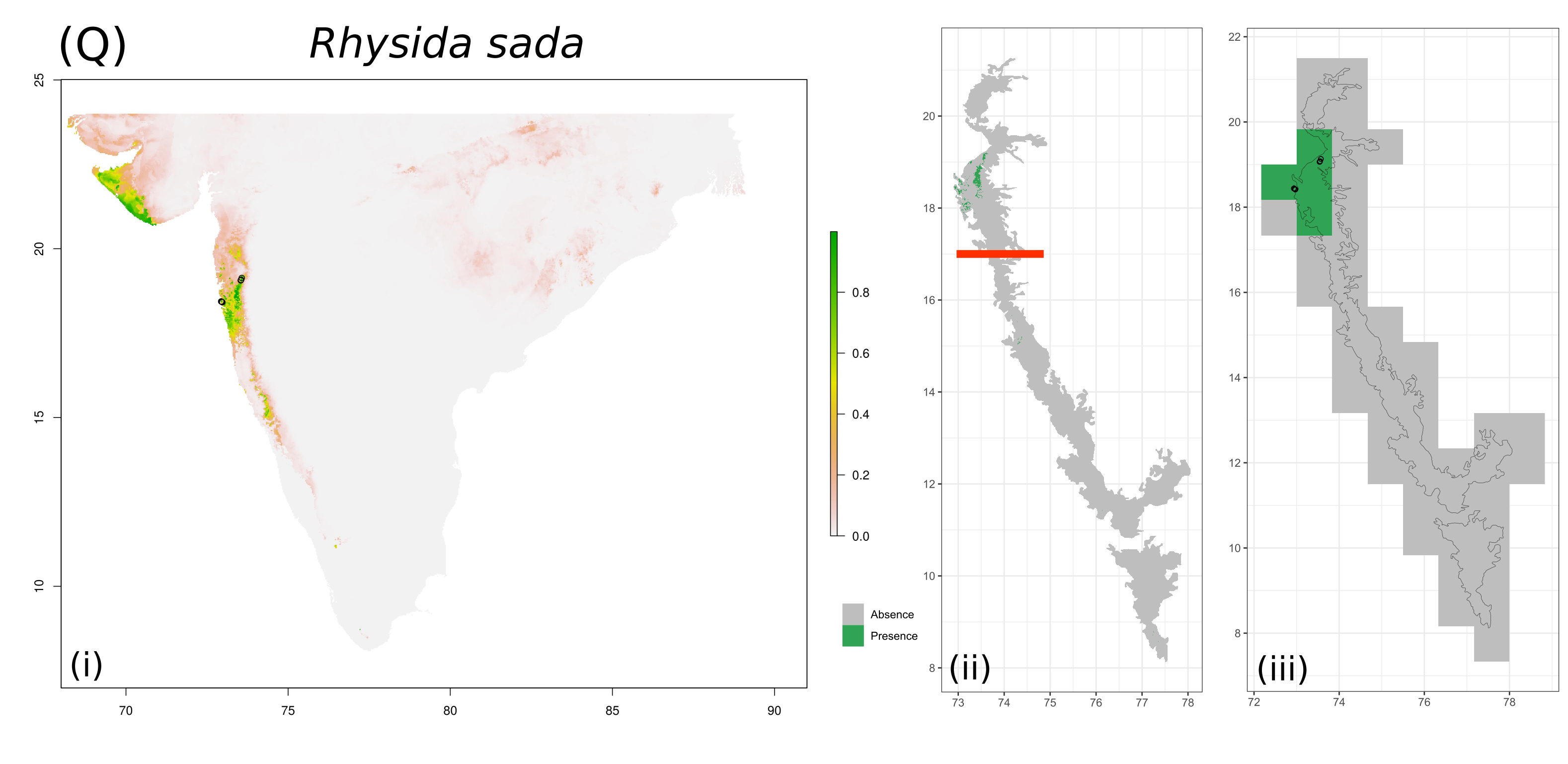

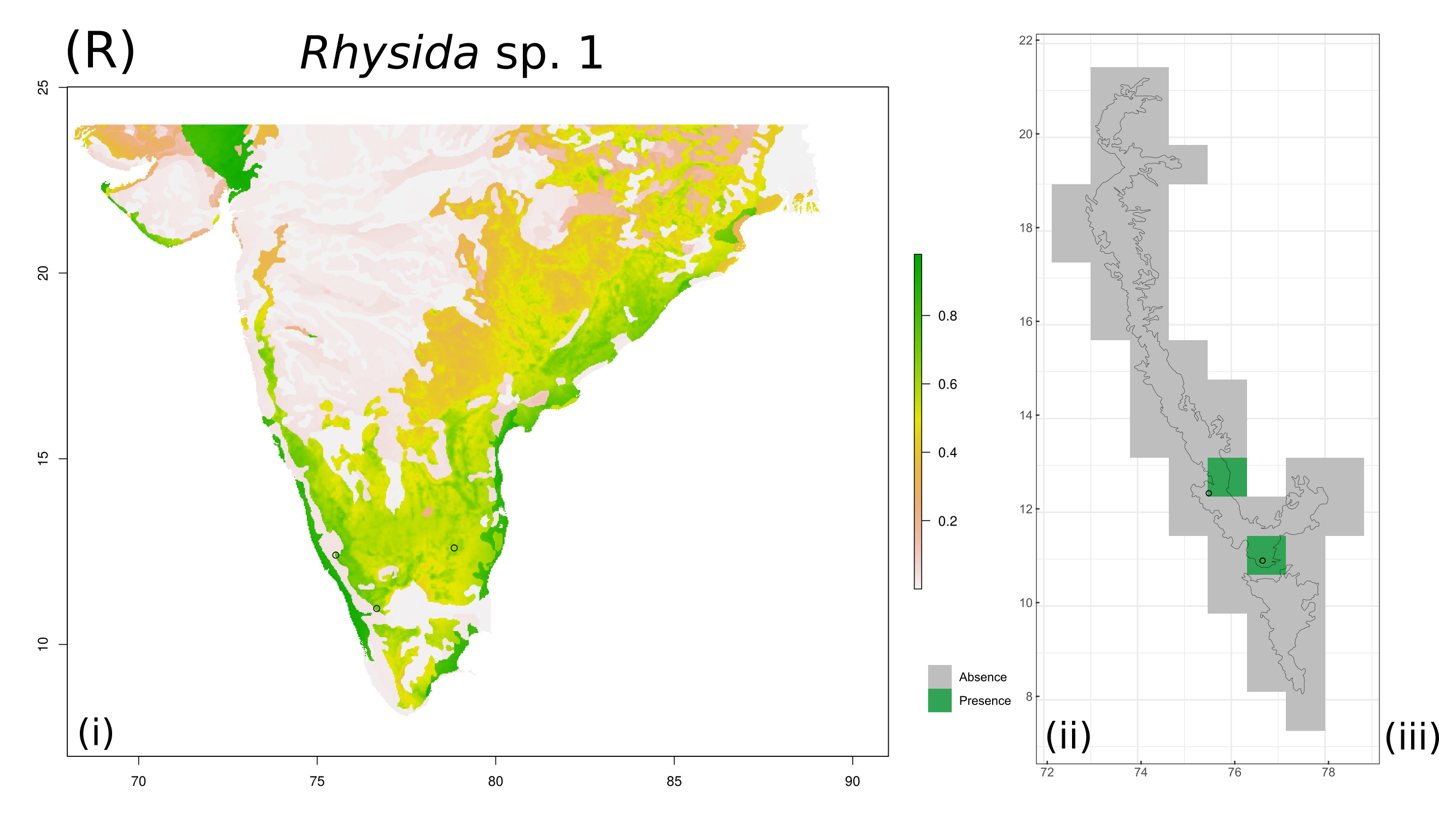

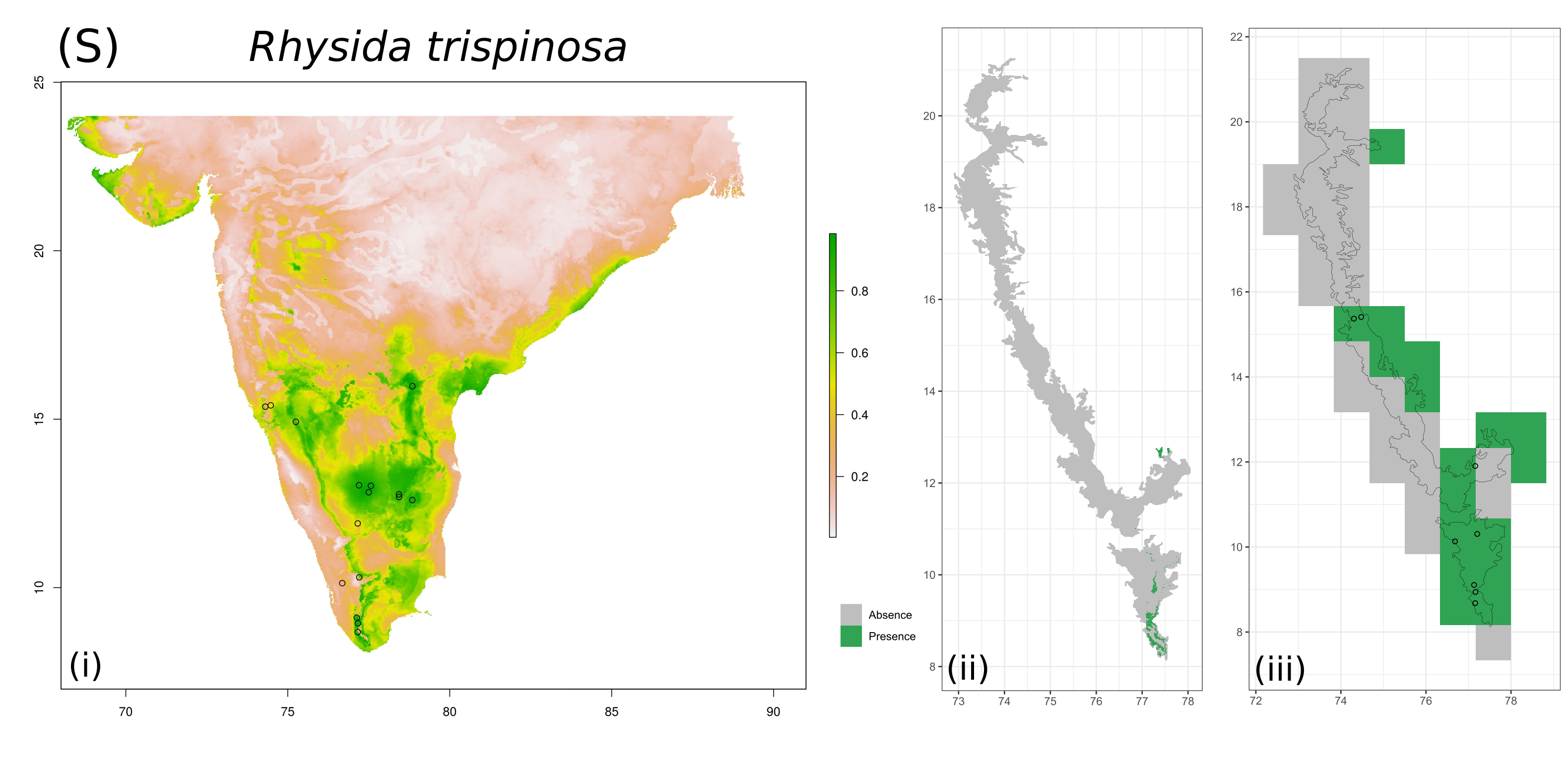

**Figure S2.3.** For each species, Maxent predictions are presented as **i.** continuous habitat suitability map for peninsular India at 0.083° ✕ 0.083° resolution from Maxent, **ii.** Presence-absence map for the Western Ghats at 0.083° ✕ 0.083° resolution obtained after applying a threshold of maximum sum of sensitivity and specificity and retaining cells above this value, **iii.** Presence-absence map for the Western Ghats aggregated to 0.83° ✕ 0.83° resolution.

For the four species listed below, the red bars in sub-figure (ii) represent limits applied to predicted range to remove small pockets of predicted habitat suitability located substantially farther away from observed presence locations:

*Digitipes coonoorensis –* removed predictions >14°N, *Ethmostigmus agasthyamalaiensis –* removed predictions >10.5°N, *Ethmostigmus coonooranus –* removed predictions >16.5°N, *Rhysida sada –* removed predictions <17°N
