## Supporting Information Appendix S3 for "Spatial patterns of phylogenetic diversity and endemism in the Western Ghats, India: a case study using ancient predatory arthropods"

**Appendix S3.** Identifying hotspots of phylogenetic endemism using the Categorical Analysis of Palaeo and Neo Endemism (CANAPE) (Mishler et al., 2014).

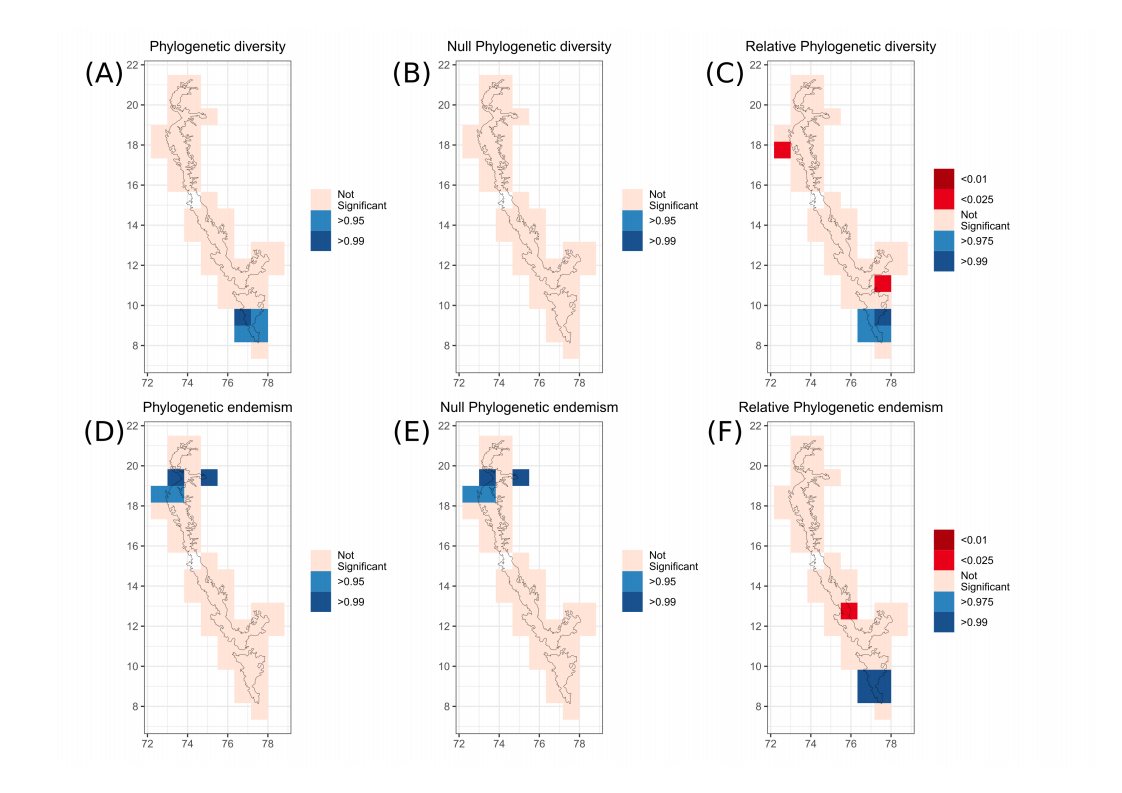

**Figure S3.4.** Significance of phylogenetic diversity and endemism indices, determined by comparison with random expectations obtained by shuffling species labels on the scolopendrid phylogenetic tree, for 0.83° ✕ 0.83° grid cells within Western Ghats, India. Cells in blue have values that are significantly greater than random expectations, cells in pink are not different from random expectations, cells in red have values that are significantly lower than random expectations. (A) phylogenetic diversity, (B) null phylogenetic diversity (calculated from phylogenetic tree with equal branch lengths – denominator of relative phylogenetic diversity), (C) relative phylogenetic diversity (phylogenetic diversity/null phylogenetic diversity), (D) phylogenetic endemism, (E) null phylogenetic endemism, (F) relative phylogenetic endemism (phylogenetic endemism/null phylogenetic endemism).

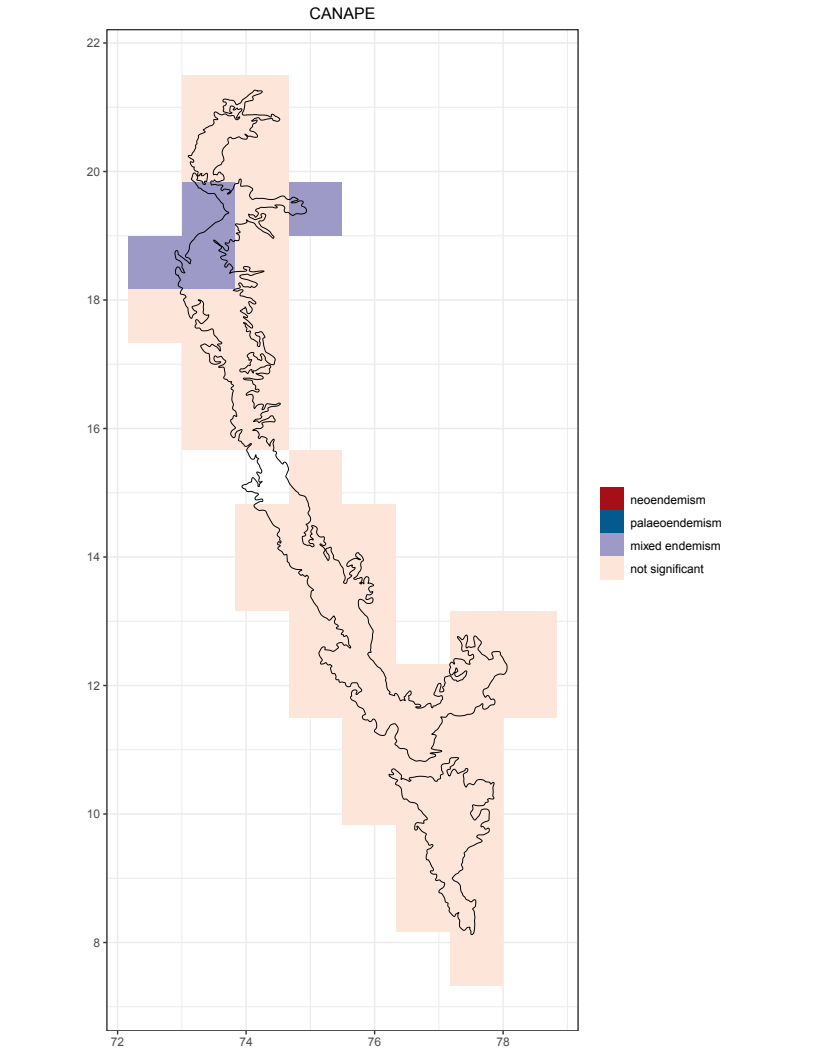

**Figure S3.5.** Classification of 0.83° ✕ 0.83° grid cells within Western Ghats, India based on CANAPE analysis (Mishler et al., 2014*). Cells in purple are areas of mixed endemism which consist of a combination of young and old endemic lineages, cells in pink are areas which did not have significant values of phylogenetic endemism.

Mishler, B. D., Knerr, N., González-Orozco, C. E., Thornhill, A. H., Laffan, S. W., & Miller, J. T. (2014). Phylogenetic measures of biodiversity and neo- and paleo-endemism in Australian *Acacia*. *Nature Communications,* *5*(4473). https://doi.org/10.1038/ncomms5473
