## Supporting Information Appendix S4 for "Spatial patterns of phylogenetic diversity and endemism in the Western Ghats, India: a case study using ancient predatory arthropods"

**Appendix S4.** Diversity and endemism measures calculated using continuous predictions of habitat suitability from Maxent species distribution model

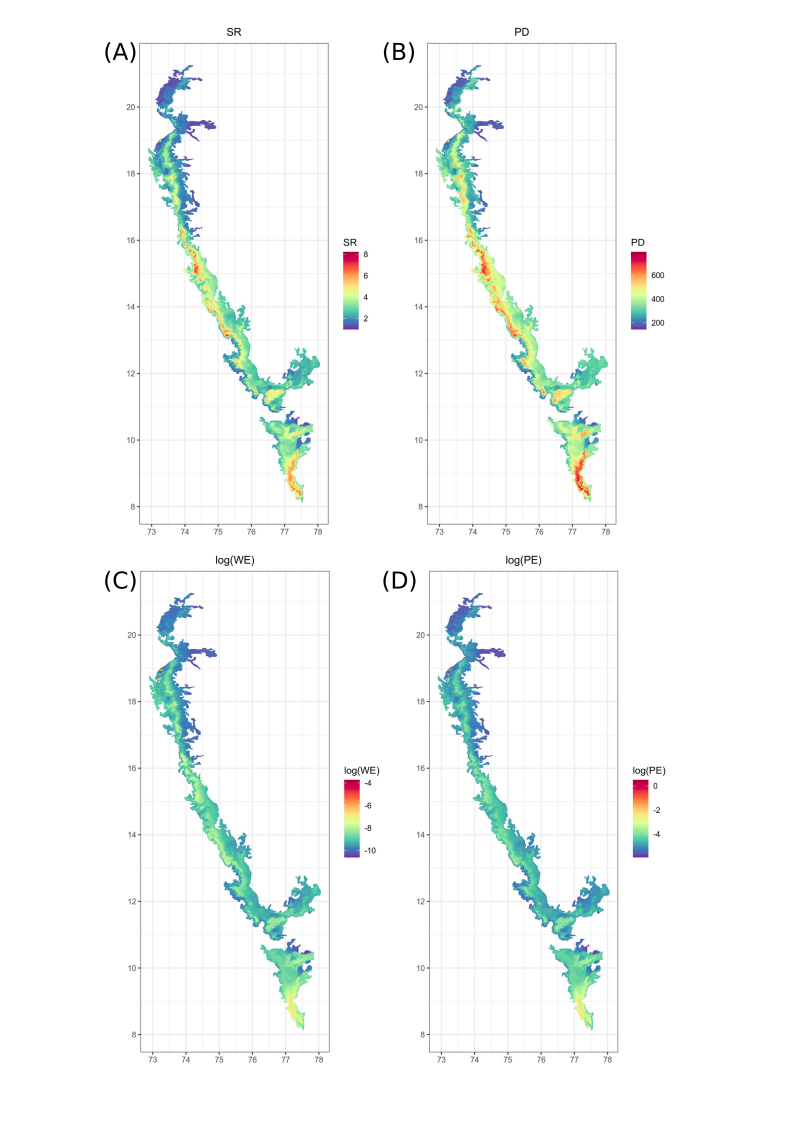

**Figure S4.6.** Maps of diversity indices for scolopendrid centipedes, (A) species richness (taxonomic), (B) phylogenetic diversity (sum of branch lengths), (C) natural log of weighted endemism, (D) natural log of phylogenetic endemism, in the Western Ghats, India in 0.083° ✕ 0.083° grid cells.
